# From sectors to speckles: The impact of long-range migration on gene surfing

**DOI:** 10.1101/803189

**Authors:** Jayson Paulose, Oskar Hallatschek

**Affiliations:** Institute for Fundamental Science and Department of Physics, University of Oregon, Eugene, OR 97403; Departments of Physics and Integrative Biology, University of California, Berkeley, CA 94720

## Abstract

Range expansions lead to distinctive patterns of genetic variation in populations, even in the absence of selection. These patterns and their genetic consequences have been well-studied for populations advancing through successive short-ranged migration events. However, most populations harbor some degree of long-range dispersal, experiencing rare yet consequential migration events over arbitrarily long distances. Although dispersal is known to strongly affect spatial genetic structure during range expansions, the resulting patterns and their impact on neutral diversity remain poorly understood. Here, we systematically study the consequences of long-range dispersal on patterns of neutral variation during range expansion in a class of dispersal models which spans the extremes of local (effectively short-ranged) and global (effectively well-mixed) migration. We find that sufficiently long-ranged dispersal leaves behind a mosaic of monoallelic patches, whose number and size are highly sensitive to the distribution of dispersal distances. We develop a coarse-grained model which connects statistical features of these spatial patterns to the evolution of neutral diversity during the range expansion. We show that growth mechanisms that appear qualitatively similar can engender vastly different outcomes for diversity: depending on the tail of the dispersal distance distribution, diversity can either be preserved (i.e. many variants survive) or lost (i.e. one variant dominates) at long times. Our results highlight the impact of spatial and migratory structure on genetic variation during processes as varied as range expansions, species invasions, epidemics, and the spread of beneficial mutations in established populations.

## I. INTRODUCTION

Range expansions have occurred in the history of many species, from plants [1] to avian [2], aquatic [3, 4], and terrestrial [5, 6] animals, including humans [7]. Over geological time scales, they have been driven by climactic changes such as glacial advance and melting in the northern hemisphere [8, 9]. More recently, anthropogenic climate change and human-mediated introduction of invasive species have driven the expansion of species into new territory [10]. These expansion events impact the genetic makeup of the population, in ways that are dramatically different from population expansions without spatial structure [11]. In particular, neutral mutations occuring during range expansions leave behind signatures that are otherwise associated with selection, such as sweeps through the population [12], allelic gradients [13], and reduction in local genetic diversity [14, 15, 16]. Understanding the patterns of neutral variation left behind by range expansions is crucial for disentangling the role of spatial structure from selection in determining genetic diversity [17].

Much of our current understanding of neutral evolution during range expansions is derived from situations where individuals migrate a short distance between generations [12, 13, 14, 15, 16, 18] In this case, the population advances through a wave of roughly constant speed separating occupied and unoccupied regions of space. Crucially, only individuals that happen to be close to the advancing front contribute to future generations, and large swathes of the population after the expansion can be traced back to a few individuals at the edge of the originating population, a phenomenon termed *gene surfing*. The resulting neutral variation shows a characteristic pattern: local diversity is strongly reduced as neutral variants segregate into uniform regions, called *sectors*, in which a single allele dominates. Nevertheless, in a radial expansion, sectors of different variants persist at long times as a result of which global diversity is maintained, as seen in Fig. 1**a–d**. These patterns persist under moderate levels of gene flow due to subsequent diffusion following the initial advance [18, 19].

**FIG. 1.**
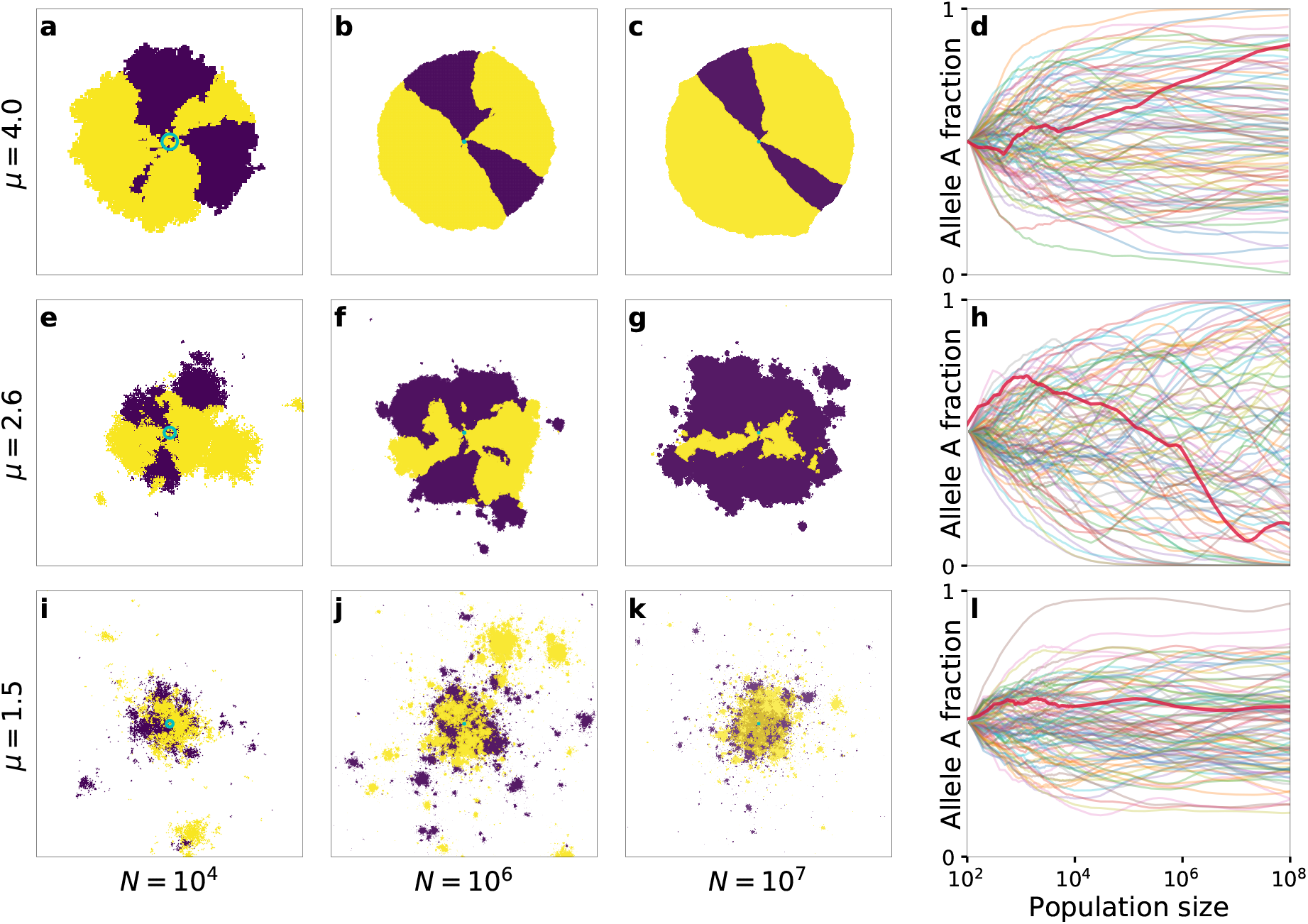
Patterns of neutral diversity under different fat-tailed dispersal kernels. **a–c**, Snapshots from a single simulation of a range expansion with kernel exponent *μ* = 4 at different population sizes *N*, starting from a homeland with *N*_0_ = 79, *q* = 2, *p*_0_ = 1/2 (cyan circle). Demes occupied by neutral alleles `A` and `B` are colored light and dark respectively. **d**, Evolution of the fraction *p* belonging to allele `A`, for 100 independent simulations (translucent curves), with the simulation from panels **a–c** shown as an opaque red curve. **e–h** and **i–l**, same as **a–c** with *μ* = 2.6 and *μ* = 1.5 respectively.

Most organisms, however, experience some amount of long-range dispersal [20, 21]. Pollen, seeds, and microorganisms are dispersed over long distances by wind and water, or by wandering or migratory animals whose excursions influence their own evolution as well. The resulting distributions of dispersal distances, also called dispersal kernels, are often “fat-tailed”: they do not have a characteristic cutoff length scale and fall off slower-than-exponentially with distance. Empirical measurements of dispersal kernels have shown that fat-tailed kernels arise in the spreading behavior of numerous species [22]. Theoretical analyses have established that fat-tailed dispersal kernels accelerate expansion dynamics [20, 23], allowing the size of the expanding population to grow faster-than-linearly with time, and strongly influence population structure by breaking up the wave of advance associated with short-ranged spreading [24, 25, 26, 27, 28, 29]

Although long-range dispersal is recognized as being consequential for range expansions [11, 30, 31, 32], its precise effects on genetic diversity are not fully understood. Whereas it is recognized that dispersal leads to monoallelic patches [24, 25, 26, 27], the conditions for patches to dominate over sectors have not been identified. Furthermore, the structural characteristics and dynamics of the patch patterns and their subsequent impact on neutral diversity have not been systematically studied. As a result, even the basic question of whether neutral variation in the originating population is maintained during dispersal-driven range expansions is unresolved. Simulation studies involving mixtures of two Gaussian (i.e. non-fat-tailed) dispersal kernels with different mean distances have shown support for contrasting effects of increasing the weight of the broader dispersal kernel on neutral diversity [33] (the so-called “embolism effect”). At low levels, founder events ahead of the expanding front of the population wipe out diversity, but at higher levels, diversity in the expanding region is maintained by serial re-introduction of variants from the interior of the population. However, other studies have argued that a reduction in diversity due to the embolism effect only occurs for dispersal along corridors and for thin-tailed dispersal kernels [34], whereas fat-tailed kernels ought to generically enhance genetic diversity [30, 34, 35, 36, 37].

Here, we study the evolution of neutral diversity in a simplified model of range expansions with dispersal events drawn from fat-tailed kernels. By analyzing a class of dispersal kernels which spans the two extremes of well-mixed growth and wavelike spreading, we obtain a comprehensive picture of neutral evolution in dispersal-accelerated range expansions. We find that long-range dispersal breaks up radial sectors into monoallelic patches, or *blobs*, but only if the kernel is sufficiently fat-tailed (Fig. 1). The characteristic size of these blobs relative to the overall size of the population can vary widely, reflecting qualitative differences in the growth dynamics for different dispersal kernels. For the broadest kernels, the spatial distribution of alleles approaches a highly fragmented speckle pattern (Fig. 1**i–l**). These patterns depart strongly from the prevailing paradigm of sectors as the spatial signature of range expansions [11, 14, 18].

We also investigate how global diversity is impacted by the breakup of sectors into blobs and speckles. By studying the growth of the typical number and size of blobs as the range expansion progresses, we show that fat-tailed kernels display the entire range of possible outcomes for neutral variation: depending on the exponent characterizing the tail of the kernel, the initial diversity can be almost perfectly preserved, or completely lost, as the expansion progresses. Strikingly, we find that longrange dispersal can in some cases erode genetic diversity compared to short-range dispersal, through a mechanism that differs fundamentally from the previously-documented embolism effect.

## II. MODEL

We consider growth into isotropic space from a compact initial population of size *N*_0_, confined to some region of space which we call the *homeland*. Each individual belongs to one of *q* allelic identities, all of which are neutral relative to each other. The population is allowed to grow into an empty range, which has a finite local carrying capacity. For concreteness, we enforce the local carrying capacity by breaking up the range into a discrete regular lattice of demes, with each able to accommodate a fixed number of individuals. We work in the limit that a newly-colonized deme reaches its carrying capacity on much shorter time scales than the time scales associated with migration. In this limit, we only need to consider two occupancy levels: demes are either completely empty or completely full, and each occupied deme has the allelic identity of the first individual that enters it. Once a deme is occupied, the local population replenishes itself constantly without changing its allelic identity, and continues to send out offspring at a fixed total rate, with migration distances *r* drawn randomly from a jump kernel *J*(*r*). The migration direction is uniformly chosen from all available directions. Although our analysis can be applied to a wide range of fat-tailed kernels, we focus on the specific form *J*(*r*) = *μr*^−(*μ*+1)^ in this work. The kernel exponent *μ* quantifies the heaviness of the tail of the dispersal kernel, with higher values corresponding to jump distributions that fall off more steeply with distance. Power-law kernels of this type encompass a broad swath of population growth dynamics, ranging from effectively well-mixed (*μ* → 0) to effectively diffusive (*μ* > *d* + 1) [38, 39]. We will henceforth refer to the population size as the number of demes, and generations are defined by the average time between migration events out of an occupied deme.

Before describing the results of multi-allele simulations, we briefly summarize the known behaviour of range expansions in power-law growth kernels for monoallelic population [38] (see Table I). The value *μ* = *d* + 1 separates two qualitatively different behaviours: for *μ* > *d*+1, the range expansion occurs via the advance of a constant-speed front separating occupied from unoccupied regions, and is similar to expansion driven by short-ranged jump kernels. For *μ* < *d* + 1, long-range dispersal events become consequential, and the radial size of the population grows faster-than-linearly with time. A second threshold at *μ* = *d* separates power-law growth of the radial size with time for *d* < *μ* < *d* + 1 from stretched-exponential growth for 0 < *μ* < *d*. As *μ* approaches zero, the population growth approaches the exponential behaviour of a well-mixed population: spatial structure becomes essentially irrelevant in this limit.

**TABLE I.** Core growth asymptotics and characteristic blob sizes. The table catalogues the asymptotic behaviour of *ℓ*(*t*) (from Ref. 38), omitting distance and time scales for *ℓ* and *t* respectively. *B*_*μ*_ ≈ 2*d* log(2)/(*μ* − *d*)^2^, and *η* = log[2*d*/(*d* + *μ*)]/log 2. Also shown is the characteristic blob size *s*\*(*r*) ≡ *ℓ*[*ℓ*^−1^(*r*)/2], introduced in the text for jump-driven growth (*μ* < *d* + 1).

| Kernel exponent | Asymptotic scaling of core radius $\ell(t)$ | Characteristic blob size, $s^*(r)$ |
| --- | --- | --- |
| $\mu < d$ | $\exp(B_\mu t^\eta)$ | $r^{\frac{\mu+d}{2d}}$ |
| $\mu = d$ | $\exp(c \log^2 t)$ | $2^{\frac{1}{4d}} e^{-\sqrt{\frac{\log 2 \log r}{d}}} r$ |
| $d < \mu < d + 1$ | $t^{1/(\mu-d)}$ | $2^{\frac{-1}{\mu-d}} r$ |
| $\mu = d + 1$ | $t \log t$ (1D) | — |
| $\mu > d + 1$ | $t$ | — |

## III. RESULTS

### A. Long-range dispersal breaks up sectors into monoallelic blobs

Fig. 1 shows snapshots from simulations of range expansions for three different kernel exponents. The narrowest kernel, *μ* = 4, corresponds to the regime of growth which asymptotically approaches a constant-speed advancing front at long times. In this regime, the population quickly coarsens into monoallelic sectors as has been well-characterized for growth due to short-ranged kernels [14, 16, 18] as seen for a representative example in Fig. 1**a–c**. At early times, boundaries between sectors of different alleles can annihilate due to the random wandering of sector boundaries from straight radial rays, but at later times, established sectors are stable against annihilation and the allelic fractions become essentially frozen in time [19, 40], up to fluctuations due to the random wandering of the boundaries between sectors. These dynamics are reflected in the evolution of allelic fractions of individual simulations, which settle to a near-constant value at long times (Fig. 1**d**).

When *μ* < *d*+1, long-ranged jumps become consequential for growth, and the resulting patterns are markedly different (Fig. 1**e–g**). Alleles still segregate into monoallelic regions, but these do not form radial sectors. Instead, the population is comprised of a mosaic of blobs of varying size, each of which has an irregular boundary but is roughly isotropic in shape. At each time point, a *core* region surrounding the homeland can be identified within which all sites are occupied. Also visible are isolated clusters of occupied sites, separated from the core by empty sites, which were colonized by the offspring of a single migrant that landed far from the bulk of the population at earlier times. These *satellite outbreaks*, visible at the outer edges of the population in each snapshot, are a characteristic feature of jump-driven growth in the presence of long-range dispersal [38, 41]. The patchiness of jump-driven range expansions arises from the continued accumulation of satellite outbreaks. The resulting disruption of sectors when *μ* < *d* + 1 will impact many known genetic consequences of range expansions, as we describe in the Discussion.

Upon comparing patterns for the two kernels displaying blobs in Fig. 1, the patterns at the broadest kernel, *μ* = 1.5 (Fig. 1**i–k**), show finer and more numerous blobs compared to the intermediate kernel with *μ* = 2.6 (Fig. 1**e–g**). Relative to the population size, blobs also appear to get finer as the range expansion progresses for *μ* = 1.5. Finally, the evolution of the allele fractions (Fig. 1**h,l**) shows different characteristics for the two kernels: each trajectory exhibits stronger variations over time for the intermediate kernel, *μ* = 2.6, and the distribution of stochastic outcomes covers a broader range of fractions. In the next section, we will connect these qualitative observations to an analysis of the typical size of blobs relative to the population size in the jump-driven growth regime.

### B. A hierarchy of doublings in time determines the characteristic size of blobs

Although the early establishment of sectors in constant-speed range expansions (such as in Fig. 1**a–c**) is stochastic, their subsequent growth over time is tied to the radial population growth and is essentially deterministic, up to random wandering of sector boundaries which becomes insignificant at long times (the growth in transverse fluctuations of the boundaries is overcome by the linear expansion of the circumference with time). By contrast, the placement and size of the monoallelic regions in the jump-driven range expansions (Fig. 1**e–k**; see Fig. 2**a** for complete time evolution of a 1D simulation) are stochastic at all stages of growth. However, the characteristic sizes of blobs incorporated into the growing core at different times (or equivalently, at different radial distances) follow a distinct pattern in the vicinity of the marginal point *μ* = *d*. This pattern was first revealed in Ref. 38, and the key features, applicable in all dimensions, are illustrated for a 1D range expansion in Fig. 2. The growth of the core radius *ℓ* as a function of time *t* from the point of seeding is constrained by a self-consistency condition: the satellites themselves grow from the accumulation of satellites at smaller scales, and must therefore follow the same growth rule *ℓ*(*t*) as the core itself (Fig. 2**b**). As a result, when the growth of *ℓ*(*t*) is faster than linear but slower than exponential (which is true in both the stretched-exponential and power-law growth regimes of Table I), the distribution of satellite clusters which join the core at time *t* is peaked at size *ℓ*(*t*/2), as shown schematically in Fig. 2**c**.

**FIG. 2.**
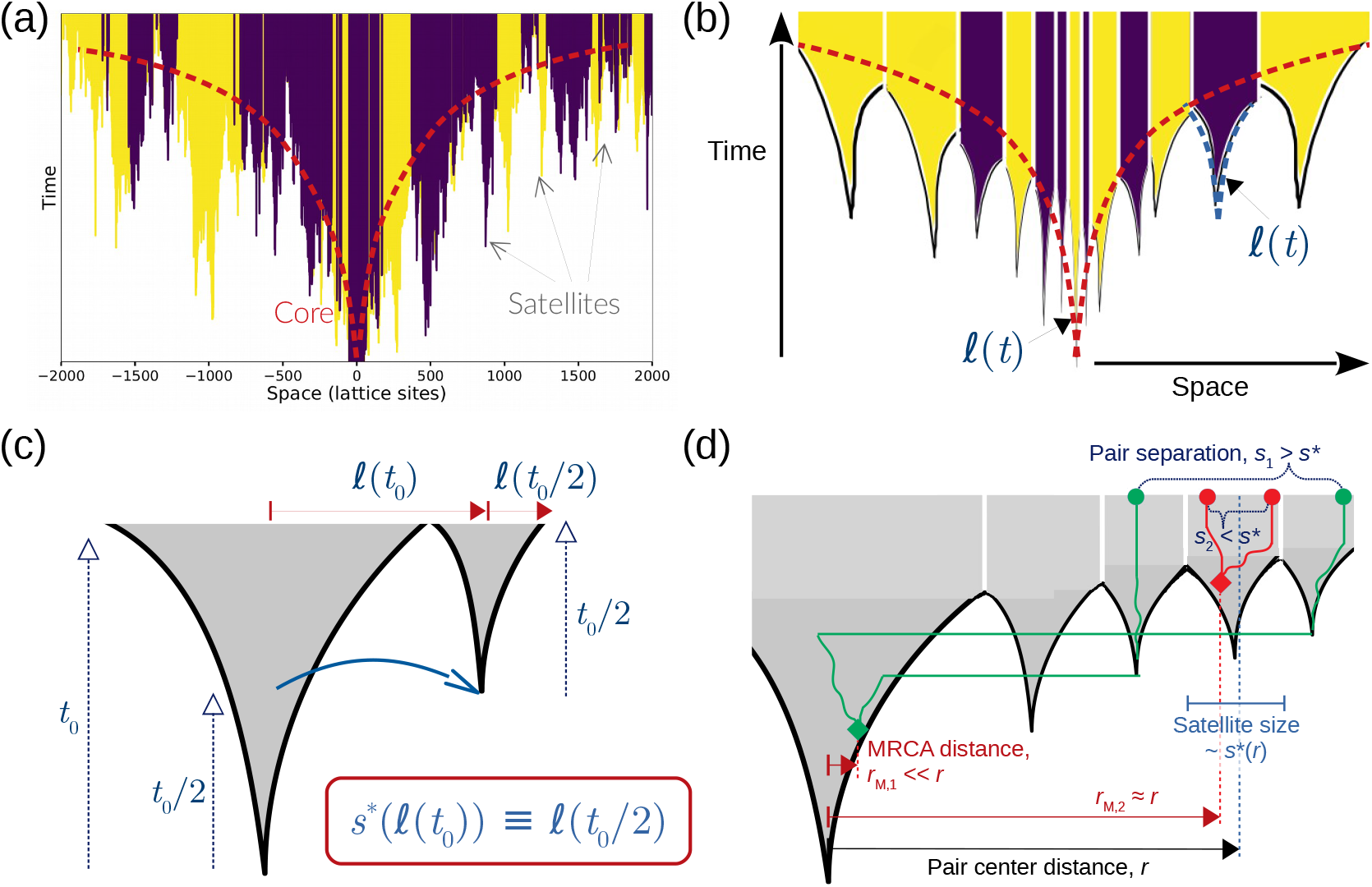
Time-doubling hierarchy for jump-driven growth and consequent ancestral pattern. **a**, Portion of a range from simulated population growth on a 1D lattice with *μ* = 1, starting from a homeland with *N*_0_ = 100 and *p*_0_ = 1/2 occupying lattice sites [−50,49]. Lattice sites are colored by their occupancy as a function of time. The dashed line indicates the coarse boundary of the region that is completely occupied at a particular time, which is termed the core region. **b**, Coarse-grained schematic of core growth due to successive mergers of satellite clusters, such as in **a**. The function *ℓ*(*t*) governing the growth of the core region must also govern the growth of the satellite clusters, which grow according to the same underlying stochastic process. **c**, For jump-driven growth under kernels with *μ* close to *d*, a self-consistency argument tying the core and satellite growth rules reveals a time-doubling hierarchy between the core and satellites at its outer edge: the typical satellite merging with the core at time *t*_0_ was seeded by a dispersal event of length *ℓ*(*t*_0_) which occurred at time of order *t*_0_/2, and gave rise to a satellite which grew for a period of order *t*_0_/2. This hierarchy defines the characteristic blob scale *s*\*(*r*). **d**, Spatial relationships between pairs of individuals (red and green discs) and their most recent common ancestor (MRCA, diamonds) reveal the time-doubling hierarchy in jump-driven expansions. Solid lines schematically trace out the genealogy of each pair to the MRCA.

The hierarchy connecting core size scales at *ℓ*(*t*) and *ℓ*(*t*/2) in jump-driven growth was developed and used in Ref. 38 to obtain the asymptotic behavior summarized in Table I, as well as the short-time approach to asymptotics. Signatures of the hierarchy were also observed in patterns left behind after parallel adaptation due to multiple mutations in a population experiencing longrange dispersal [41]. In the context of neutral diversity in range expansions, this hierarchy links the sizes *s*(*r*) of monoallelic regions to their distance *r* from the center of the range expansion: we expect *s*(*r*) ~ *ℓ*[*ℓ*^−1^(*r*)/2]. The asymptotic growth forms (Table I) predict qualitatively different relationships for the size of the largest satellite clusters in the population relative to the core population itself in the various growth regimes. When *μ* < *d*, we expect *s*(*r*)/*r* ~ *r*^(*μ*−*d*)/(2*d*)^, a decreasing function of *r*: the characteristic size of the largest clusters shrinks relative to the core size itself as the range expansion advances. By contrast, when *d* < *μ* < *d* + 1, we expect *s*(*r*)/*r* ~ 2^−1/(*μ*−*d*)^, a constant as the core size increases. The representative patterns for *μ* = 2.6 and *μ* = 1.5 in Fig. 1 are consistent with these qualitative features. When *μ* = 2.6, the characteristic sizes of the blobs at the edge of the core do not vary significantly relative to the core size, whereas for *μ* = 1.5 the blobs appear smaller relative to the core as the simulation progresses.

To quantitatively test the hierarchy of blob-core sizes predicted above, we measured spatial relationships among pairs of individuals and their most recent common ancestor (MRCA). As Fig. 2**d** illustrates, the time-doubling hierarchy predicts that a pair of individuals centered at distance *r* is likely to belong to the same satellite if their separation is much less than *s*(*r*), and hence have an MRCA located at a distance of order *r* from the origin. By contrast, individuals separated by distances much larger than *s*(*r*) are likely to belong to independently growing satellites which were seeded by different long jumps from the core; their MRCA is located at distances much closer to the origin. In the full evolution, the radial position *r*_M_ of the MRCA of a pair of occupied sites separated by a distance *s* about a distance *r* from the origin is a stochastic variable. However, we expect the *average* of the MRCA positions of many such pairs to fall from *r* to 0, over a separation distance of order *s*\*(*r*) ≡ *ℓ*[*ℓ*^−1^(*r*)/2].

Measurements of pair-MRCA relationships in 1D simulations confirm these expectations (Fig. 3; see Methods for details). In all growth regimes, the average MRCA position *r̅*_M_ falls with separation *s* over a characteristic decay scale which grows with increasing center-pair distance *r* from the origin, consistent with satellite sizes growing larger as the range expansion progresses (Fig. 3**a–c**). Upon rescaling the pair separation with the proposed blob scale *s*\*(*r*), data for different values of *r* collapse onto curves that depend only on the jump kernel (Fig. 3**d**), with a sharp fall in mean MRCA distance when *s/s*\* ~ 1. The data collapse reflects the relevance of the time-doubling hierarchy in setting the characteristic scales underlying the stochastic blob size distributions during the range expansion. Note that merely rescaling pair separation with the center-pair distance does not lead to collapse of the data curves in the stretched-exponential and marginal growth cases (insets to Fig. 3**a–b**), since the characteristic blob size falls relative to the size of the entire population as the expansion proceeds. By contrast, for power-law growth the ratio *s*\*(*r*)/*r* is independent of *r*, so rescaling *s* with *r* also causes the data curves to coincide (Fig. 3**c**, inset).

**FIG. 3.**
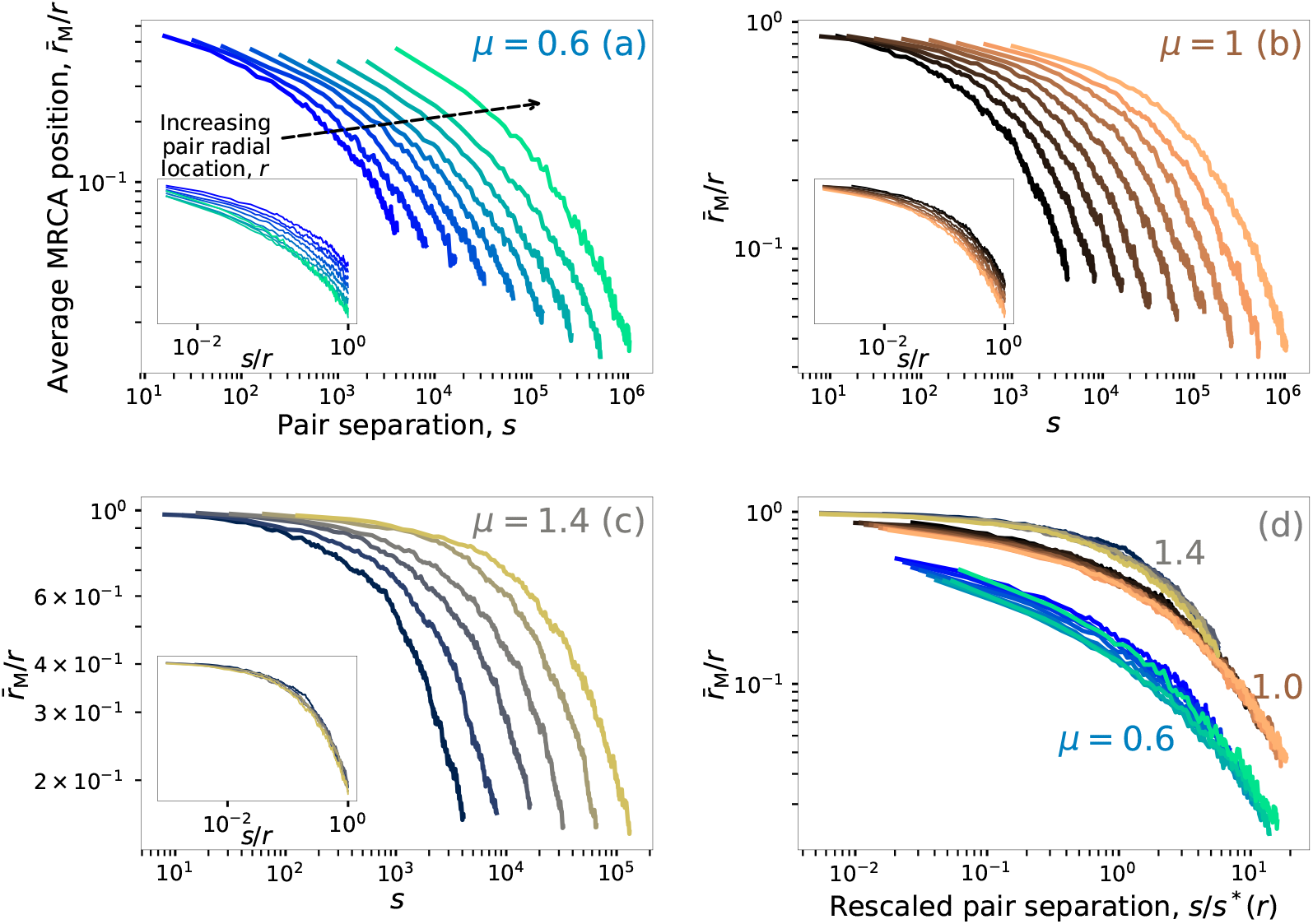
The time-doubling hierarchy is revealed in MRCA positions of pairs of sites in 1D simulations. **a–c**, Dependence of the average MRCA radial position *r̅*_M_ on the separation *s* between the pair in outbreaks starting from a single individual at the origin, with each curve corresponding to pairs whose mean position lies at a center-pair distance *r* from the origin. In all panels the leftmost curve corresponds to *r* = 4096 and subsequent *r* values are related to the leftmost curve by successive multiples of two (*r* = 8192, 16384, …, 1048576). Values of *r/r̅*_m_ close to one signify that the pair belongs to the same satellite, whereas pairs split across different originating satellites have *r/r̅*_M_ ≪ 1. Panels **a**, **b** and **c** correspond to stretched-exponential (*μ* = 0.6), marginal (*μ* = 1) and power-law (*μ* = 1.5) growth respectively. The inset shows the same curves, with the pair separation rescaled by the radial position. **d**, curves from **a–c** rescaled by the blob size scale *s*\*(*r*) = *ℓ*[*ℓ*^−1^(*r*)/2] for each value of *r*.

### C. A coarse-grained model of blob replication predicts distinct outcomes for neutral diversity in different growth regimes

Next, we investigate the effect of the time-doubling hierarchy of blob sizes on the global neutral genetic diversity as the range expansion progresses. Since each satellite originated from a single founder, the isolated growth of satellites acts as a coarsening mechanism which locally reduces diversity. However, all individuals in the core can contribute long-distance migration events, so the seeding of new satellites provides a mechanism to maintain global genetic diversity in the population. The competition between coarsening and diversification determines the fate of neutral diversity during jump-driven growth.

We now develop a semi-deterministic model for the evolution of the average neutral heterozygosity in jump-driven range-expansions, which combines the deterministic placement and growth of satellite domains with random draws of the allelic identity of each domain. The jump-driven growth dynamic has two consequences for the dynamics of seeding and coarsening: i. The allelic identity of a typical satellite domain joining the core at time *t* is determined by a seed drawn from the gene pool of the core at time *t*/2; ii. The seed contributes its allelic identity to the entire satellite with size of order *ℓ*(*t*/2), which sets the scale of the coarsening at time *t*. These facts suggest that the evolution of heterozygosity is best described over “generations” that involve doublings in *time*, not population size. Specifically, in the deterministic approximation, the state of the core at time *t* provides sufficient information to generate the core at time 2*t*. The sizes of the satellites added to the population between *t* and 2*t* is set by geometry and the form of *ℓ*(*t*) in the deterministic approximation, but the random drawing of seeds from the core introduces stochasticity to the process. This hierarchy can be formalized in an urn-like model of core growth through the accumulation of satellites, which produces stochastic outcomes which can be numerically studied, see Appendix A.

Analytical progress can be made through additional simplifying assumptions. First, we ignore the spatial structure of the core between doublings. Instead, we treat the core at each doubling as a spatially homogeneous mix of alleles from which seeds are randomly drawn. Second, rather than attempting to capture the time-evolution of all alleles, we track the global heterozygosity *H*, a commonly-used metric of population-level genetic variation. The heterozygosity is defined as the probability that a random pair sampled from the population will have alleles of different identity. To be concrete, we consider a doubling which evolves a homogeneous core of radial size *ℓ*(*t*_0_) to size *ℓ*(2*t*_0_). In the deterministic approximation, the doubling requires the addition of *g*_1_ satellites, where *g*_1_ ~ [*ℓ*(2*t*_0_)/*ℓ*(*t*_0_)]^*d*^ is the number of patches of the *t*_0_-population which fit into the core at time 2*t*_0_ (Fig. 2**d**). The allelic identities of the new satellites are stochastically determined by random draws from the population within *ℓ*(*t*_0_). The heterozygosity *H*_1_ of the new population is therefore a stochastic random variable, but its expected value can be related to the initial heterozygosity *H*_0_ by evaluating the probability that a random draw of two individuals from the new population is biallelic. This probability is zero if both individuals are drawn from the same new satellite, and *H*_0_ in all other situations. Ignoring the variation in size of the new satellites (see Appendix A for more details), the expected heterozygosity after one doubling is

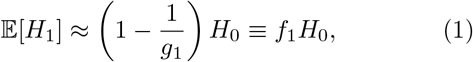

where *i* ∈ {1, 2…, *g*_1_} indexes the new satellites.

Equation (1) captures the balance between coarsening and seeding in the deterministic approximation to jump-driven growth, with the influence of the growth dynamic manifested in the factor *g*_1_. The value *f*_1_ is always less than one because of the coarsening due to monoallelic satellites; however, it can be close to one if many small satellites are seeded in each doubling, so that *g*_1_ ≫ 1 for a large number of satellites. Upon comparing Eq. (1) with the evolution of heterozygosity *H*_1_ = (1 − 1/*N*)*H*_0_ for one generation of a haploid Wright-Fisher model with population size *N*, we may interpret *g*_1_ as the *effective population size* determining the strength of genetic drift during the first “generation” which takes the population from time *t*_0_ to 2*t*_0_.

To connect the evolution described above to the growth of a population out of a spatially homogeneous, compact homeland of population size *N*_0_, we interpret the homeland as a core which has grown to a radius *ℓ*_0_ ≡ (*N*_0_/*ω*_*d*_)^1/*d*^, where *ω*_*d*_ is the volume of the unit *d*-dimensional sphere (*ω*_1_ = 2, *ω*_2_ = *π*). The growth rule *ℓ*(*t*) can be inverted to define a characteristic time *t*_0_ ≡ *ℓ*^−1^(*ℓ*_0_) which sets the time scale over which heterozygosity is tracked in the coarse-grained model. The evolution of the average heterozygosity for growth over times *t* ≫ *t*_0_ is given by the cumulative effect of *n* ≈ log_2_(1+*t/t*_0_) successive doublings, each indexed by *m*:

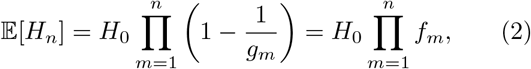

where *g*_*m*_ is the number of satellites generated in the *m*th doubling. Over long times, therefore, the neutral diversity in the effective population of satellites is determined by the behavior of the product 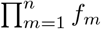, which in turn is determined purely by geometry and the form of the growth rule *ℓ*(*t*).

We now investigate the evolution of the average heterozygosity predicted by Eq. (2) for the asymptotic growth forms of Table I. For kernels with *μ* < *d*, the asymptotic growth rule is *ℓ*(*t*) ∝ exp(*B*_*μ*_*t*^*η*^), where *η* = log_2_[2*d*/(*d* + *μ*)] approaches one as *μ* → 0 (the well-mixed limit) and zero as *μ* → 1. For successive doublings starting from a homeland of size *ℓ*_0_ = *ℓ*(*t*_0_), we find 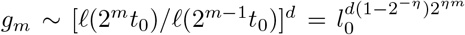. Away from *η* ≈ 0 (i.e., from *μ* → *d*), the number of satellites per doublings grows extremely fast 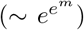 with the number of doublings. As a result, *f*_*m*_ ≈ 1 − 1/*g*_*m*_ quickly approaches one with increasing *m*, leading to fast convergence of the product in Eq. (2) to a constant value (see Appendix A for details). The coarse-grained model therefore predicts that neutral diversity can be preserved at long times in this model: complete fixation of one allele is avoided at long times, and a finite average heterozygosity is reached within a few doublings.

In contrast to the stretched-exponential growth rule, a power-law growth rule *ℓ*(*t*) ∝ *t*^1/(*μ*−*d*)^ has no intrinsic size scale: the relative geometry of the new satellites added during a deterministic doubling is independent of the size of the core. As a consequence, *f*_1_ is independent of the size of the core, and *f*_*m*_ = *f*_1_ ≡ *f* remains *constant* during subsequent doublings. Eq. (2) then predicts that the heterozygosity *H*_*n*_ = *f*^*n*^*H*_0_ decays exponentially with doublings. Unlike the stretched-exponential growth regime, the effective population of monoallelic satellites does not grow fast enough to evade fixation of a single allele over many doubling “generations”. The effective population of satellites loses genetic diversity in a process akin to genetic drift, due to the existence of a constant “effective population size” *N*_e_ ~ *g* ~ [*ℓ*(2*t*)/*ℓ*(*t*)]^*d*^ = 2^*d*/(*μ*−*d*)^. We emphasize that the lapse of one generation for the effective population of satellites corresponds to a *doubling* of time, in contrast to a fixed increment of time for ordinary genetic drift.

Kernels with *μ* = *d* represent a marginal situation between power-law and stretched exponential growth (Table I), with a core growth *ℓ*(*t*) ~ exp(log^2^*t*) that is faster than any power law yet slower than stretched-exponential growth at long times. Repeating the above analysis for growth out of a homeland of radius *ℓ*(*t*_0_), we find that the number of satellites, and hence the effective population size, grow *exponentially* in the number of doublings *m* as 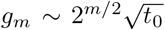. This growth is fast enough for the average neutral heterozygosity, (2), to converge to a finite value which lies between *H*_0_ and zero (see Appendix B). In contrast with the power-law growth regime, the growth of the effective population size allows neutral diversity to be partially preserved over long times; however, the growth of *g*_*m*_ with doublings, and hence the convergence of the average heterozygosity, are significantly slower than for the stretched-exponential regime.

In summary, the coarse-grained, semi-deterministic approximation of the jump-driven growth provides a minimal model that allows us to evaluate the competing effects of coarsening and diversification during jump-driven range expansions. The model predicts that that when *μ* ≤ *d*, the diversifying effect dominates and average heterozygosity converges to a finite value at long times. By contrast, the coarsening mechanism dominates when *d* < *μ* < *d* + 1, and diversity is steadily lost over time in a process similar to genetic drift in constant-size populations.

### D. Heterozygosity evolution in simulations is consistent with the coarse-grained model

To test whether the predictions of the coarse-grained model hold under the full dynamics, we investigate the evolution of neutral diversity in simulations with *q* = 2, for which the global diversity is given by *H* = 2*p*(1 − *p*) where *p* is the fraction of one allele, starting from its initial value *p*_0_ = 1/2. For each kernel, hundreds of independent simulations were performed and their heterozygosities were averaged to produce the ensemble-averaged heterozygosity 〈*H*〉. Measurements of 〈*H*〉 from both 2D and 1D simulations are reported in Fig. 4.

**FIG. 4.**
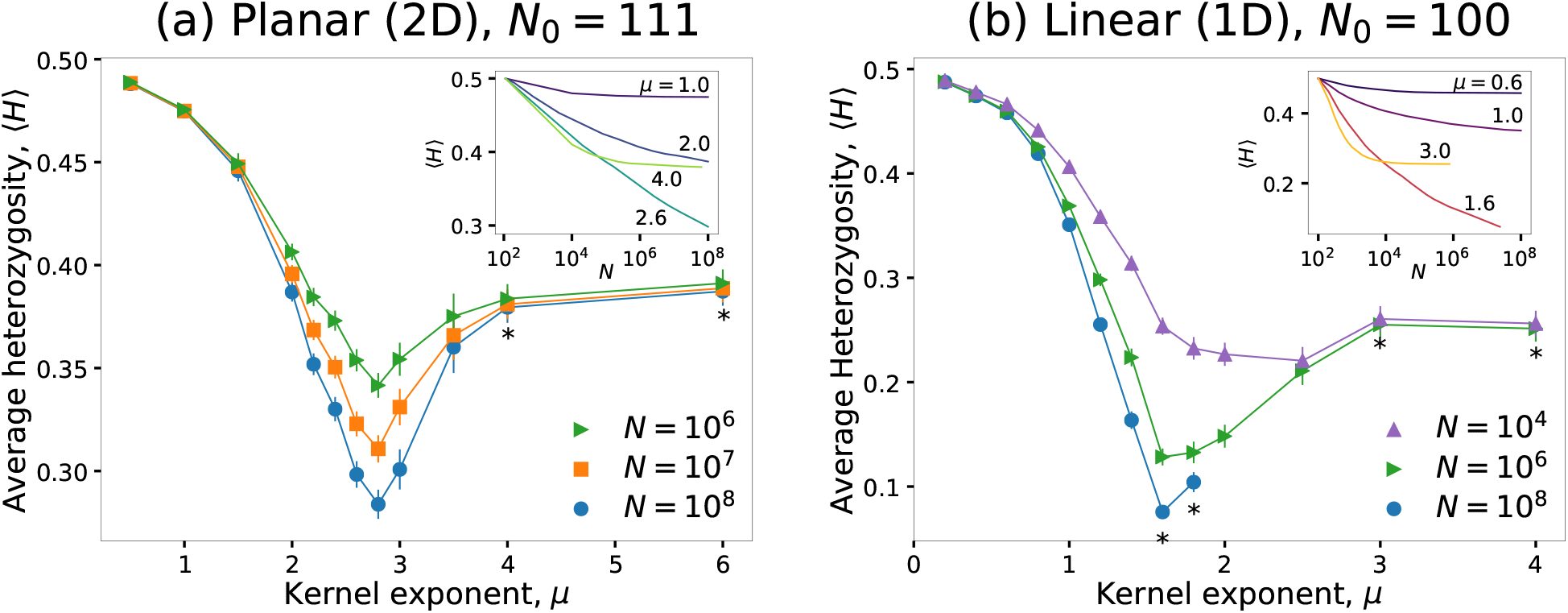
Average final heterozygosity is non-monotonic in kernel exponent. Average heterozygosity measured from simulations at different values of kernel exponent *μ* for 2D (**a**) and 1D (**b**). The initial homeland size is *N*_0_ = 111 in **a** and *N*_0_ = 100 in **b**. The average is shown for three values of final population size. Error bars show standard error of the mean across 200–800 independent simulations. The insets show the ensemble-averaged heterozygosity traces against population size for a subset of the simulated kernels (labeled). Asterisks denote data points for which the largest system size was restricted to the following values due to simulation constraints: (2D) *N* = 6.4 × 10^7^ for *μ* = 4 and *N* = 5.6 × 10^7^ for *μ* = 6; (1D) *N* = 2.4 × 10^7^ (*μ* = 1.6), *N* = 7 × 10^6^ (*μ* = 1.8), *N* = 8 × 10^5^ (*μ* = 3), *N* = 6 × 10^5^ (*μ* = 4).

For both planar and linear range expansions, the measured average heterozygosities are consistent with the predicted trends. In 2D simulations (Fig. 4**a**), the average heterozygosity does not change significantly between final population sizes of 10^7^ and 10^8^ for *μ* < 2 and *μ* > 3, signifying convergence to a finite value (see inset and Fig. A7 for evolution of heterozygosities with population size). However, in the range of kernel exponents 2 ≤ *μ* ≤ 3, the average heterozygosity continues to drop with population growth over the entire range of simulated sizes, which is consistent with a steady loss of diversity as the population grows for this range of kernels. The results of 1D simulations (Fig. 4**b** and Fig. A8) are qualitatively similar, but with persistent loss of diversity occuring for kernel exponent values 1 ≤ *μ* ≤ 2. These results support the conclusion that coarsening leads to loss of neutral variation when *d < μ < d* + 1, whereas diversification preserves neutral variation for *μ* < *d*. The preservation of partial diversity when *μ* > *d* + 1 is due to the known mechanism of sector formation, since the range expansion occurs via a constant-speed front in this range.

Although the qualitative difference between kernels with *μ* < *d* and *μ* > *d* is readily observed, the outcome of the marginal situation *μ* = *d* is less clear. In both 2D and 1D, the average heterozygosity is still falling up to the largest simulated population sizes (Fig. 4). However, this observation is consistent with the conclusions from the coarse-grained model. For marginal growth, the convergence of the product in Eq. (2) occurs over many time-doubling “generations”. In real time, this implies that the population must grow over many orders of magnitude before the average heterozygosity converges to its limiting value. In Fig. A3, we show that the heterozygosity decay observed in simulations is consistent with Eq. (2), for several homeland sizes in both 1D and 2D. According to our theory, the largest simulations we have run correspond to roughly seven time-doubling generations, whereas convergence is expected after 15-20 doublings. Therefore, our simulated population sizes are still many orders of magnitude too small for convergence to to be observed.

Appendix B also reports additional predictions for the average heterozygosity from the coarse-grained model in the stretched-exponential (*μ* < *d*) and power-law (*d < μ < d* + 1) regimes. Comparisons of these predictions with simulation results (Fig. A2 and Fig. A4) demonstrate that the coarse-grained model, despite its simplifications and approximations, reproduces many features of the heterozygosity trends for kernels across different regimes.

## IV DISCUSSION

We have demonstrated that long-range dispersal can dramatically impact the local structures and global trends of neutral genetic diversity left behind by a range expansion. Specifically, dispersal kernels with a power-law tail characterized by exponent *μ* < *d* + 1 show patterns of diversity that qualitatively differ from the radial sectors left behind by populations which expand through local dispersal (cf. Fig. 1). Locally, regions dominated by a single allele form nearly-isotropic patches, termed “blobs”, whose typical size increases as the range expansion progresses. Upon considering the growth in blob size relative to the size of the population itself, two distinct regimes have been identified. When *d* < *μ* < *d* + 1, the typical size of the largest blobs is a constant kernel-dependent fraction of the size of the population (i.e. *s*\*(*r*)/*r* = const., Table I). By contrast, when *μ ≤ d*, blob sizes shrink relative to the population as the expansion progresses, leading to a pattern of fine monoallelic speckles (Fig. 1**i–k**).

As a consequence of a time-doubling hierarchy inherent to jump-driven growth, we have identified an “effective population size” of blobs generated during a doubling in time. The evolution of this effective population over “time-doubling generations” (which are constant increments in *log* time) follows distinct trends in the two regimes of jump-driven growth: the effective population size is stagnant in the first growth regime (*d < μ < d*+1), but grows in the second regime (*μ ≤ d*). As shown schematically in Fig. 5, the resulting stochastic fluctuations in allele fractions accumulate to produce very different outcomes from superficially similar growth mechanisms in a process akin to genetic drift, but where each generation is an increment in log_2_(*t/t*_0_) rather than a fixed time interval. Although the simulations reported above only include two distinct alleles, our coarse-grained model applies for generic initial allele numbers and frequencies, and our predictions for the evolution of heterozygosity are independent of the number of distinct alleles present (see Appendix C and Fig. A6).

**FIG. 5.**
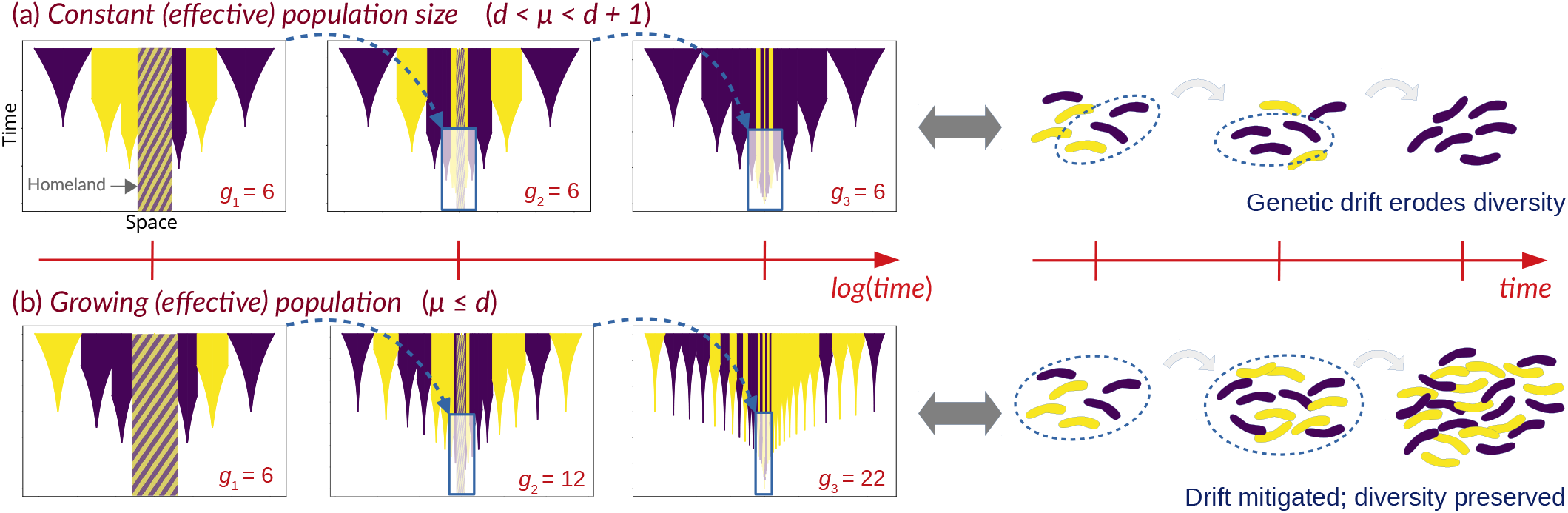
The effective population of satellites determines the fate of neutral diversity. The coarse-grained model of jump-driven growth treats a range expansion out of a well-mixed homeland as a succession of time-doublings indexed by *i*, each of which adds *g*_*i*_ monoallelic satellites drawn at random from the population after the previous doubling. The *gi* values are determined by the core growth rule, *ℓ*(*t*) (Table I). **a**, In the power-law growth regime, *g*_*i*_ remains constant over successive time-doubling “generations” (left), where the lapse of a generation in the effective population requires time to double in the actual population (i.e., generations are increments in log_2_(*t*)). As a result, genetic diversity is eroded over time due to chance events, in a process analogous to genetic drift in a well-mixed population with the number of individuals kept constant (right). **b**, By contrast, in the marginal and stretched-exponential growth regimes, the effective population of satellites grows with each time-doubling generation, and the growth is fast enough to mitigate drift.

The breakup of sectors into blobs and speckles has important genetic consequences. As can be seen in Fig. 1**a–c**, the sector geometry generated by constant-speed range expansions leaves a signature of the direction of expansion on local genetic patterns [14, 18], which can be detected in principal component maps of genetic variation [42]. By contrast, the relation between local blob geometry and global population history in jump-driven expansions (Fig. 1**e–k**) is more subtle: individual blob shapes are isotropic and do not directly reveal the expansion direction. Blobs do get larger with increasing distance from the homeland on average, but the significant stochastic variation in blob sizes makes this signature difficult to detect. Since neighboring blobs could have been seeded by migrants from well-separated regions, jump-driven range expansions are also expected to have significantly higher levels of mixing compared to sectored expansions. This mixing reduces the positional advantage conferred to mutations which arise near the edge of the expanding population, thus mitigating the gene surfing effect [12, 13]. The finer structure and increased mixing due to blobs are likely to have an impact on the evolutionary effects of geographic structure such as reduced adaptive potential [43], response to inbreeding depression [44], and expansion load [45].

Whereas we have focused on describing the expansion of a population into previously unoccupied territory, our results are applicable to other biological expansions as well. The growth dynamics of our model also applies to the spread of beneficial mutations from a localized region into an established wildtype population with a spatially uniform population density [19]. In this context, our model would describe the patterns of variation caused by the spread of distinct beneficial mutations with similar fitness effects — a *soft sweep* [41, 46, 47] — out of a small region (the homeland) which experienced a selection pressure earlier than the rest of the population. Analogues of the patterns and mechanisms described here could also play a role in within-host viral dynamics during infections [48, 49] and in cancer metastasis [50, 51].

An outstanding question regarding the population-genetic consequences of dispersal on range expansions has been whether enhanced dispersal preserves or erodes diversity relative to short-range migration. In a previous study of mixtures of two non-fat-tailed kernels with dif ferent characteristic jump lengths, intermediate levels of longer-range jumps were shown to reduce neutral diversity relative to low or high levels in narrow corridors [33]. The reduction occurred when a fortuitously-placed satellite from a pioneer seed quickly filled the width of the corridor and blocked other alleles from advancing, a mechanism termed the embolism effect (this is a version of the more generic “founder takes all” mechanism [52]). However, restriction of growth along a narrow corridor is essential for the embolism effect to wipe out genetic variation [34]; in a radial expansion, embolisms would only suppress diversity within certain angular ranges, and neutral variation would persist in the form of sectors at long times. Furthermore, the embolism effect was demonstrated for kernels with a strict upper limit to the allowed jump distances, which restricted the ability of individuals from the interior of the population to contribute to diversification. For these reasons, other studies have speculated that fat-tailed kernels without an upper cutoff in dispersal distance might not experience the embolism effect, and might enhance diversity relative to short-ranged dispersal in all cases [34, 35, 36, 37].

In the context of these previous studies, a key result of our work is that even fat-tailed kernels without a cut-off can induce a loss of neutral variation. Similarly to Ref. 33, we have shown that boosting long-range dispersal has a non-monotonic effect on diversity: intermediate kernels (i.e. with power-law exponent *d* < *μ* < *d* + 1) preserve less variation than broader (*μ* < *d*) or narrower (*μ* > *d* + 1) kernels. However, the loss of neutral variation we observe for intermediate kernels (*d < μ < d* + 1) is fundamentally different from the embolism effect: it relies on the fat-tailed nature of the kernel, which allows a finite probability of jumps that span the whole population regardless of its size. Through a mechanism reminiscent of genetic drift, the expanding population is engulfed by satellites belonging to a single allele purely through chance. As a result, the decay of heterozygosity is observed even in 2D radial range expansions which are not confined to a corridor. Even in a linear geometry, our mechanism can be distinguished from the embolism effect. For a 1D expansion from a central source growing out in two directions, the embolism effect could support two different alleles growing out in the left and the right directions. By contrast, in our model diversity is completely eroded when 1 < *μ* < 2 in 1D, with only *one* allele surviving at very long times.

Theoretical and computational studies (e.g. Refs. 12, 13, 15, 33, 34, 35, 36, and 37) and laboratory experiments on model organisms (Ref. 14), although highly highly simplified relative to real-world populations, have nevertheless provided heuristics and mechanisms that have guided the interpretation of field genomic data. In the context of range expansions, the concepts of gene surfing, sectors, and embolisms have been invoked to explain patterns of genetic variation in a wide range of plant [53, 54, 55], animal [56, 57], and microbial [58] populations. Our work shows that these heuristics are incomplete when long-range dispersal is present: sectors give way to *blobs* and *speckles* with increasing levels of dispersal; the new mechanism of *engulfment* can engender diversity loss even for fat-tailed kernels in the absence of corridors, where the embolism effect does not apply.

Many interesting avenues for further study can be identified. By filling up demes instantaneously and irreversibly, we have focused in this work on the genetic patterns seen immediately after colonization. We have ignored subsequent reshuffling of alleles among demes, which would blur the boundaries between blobs at later times and smear out the predicted spatial patterns over time. This smearing would impact the ability to detect the patchiness of jump-driven range expansions at later times, with the problem being more severe closer to the homeland. While previous studies have shown that gene segregation due to the range expansion persists for appreciable periods of time beyond the colonization for both short-range [18] and long-range [34] dispersal, a more refined spatial model incorporating the exchange of individuals among demes of finite population size would address the question of how long the patterns remain measurable at different distances from the homeland. (Note that the blurring of patterns near the interior during the range expansion does not impact the evolution of global diversity, since the spatial organization of alleles in the core is irrelevant to the identity of satellites generated in the time-doubling hierarchy.) Such a model could also be used to study the interplay of genetic drift within demes and the large-scale diversity evolution captured in our coarse-grained model. In addition, the applicability of our results to continuous populations without a deme structure could be studied by introducing long-range dispersal into continuum population genetics models [59, 60] and simulations [61].

Our results show that spatial constraints fundamentally alter the mathematical structures underlying neutral evolution in expanding populations. Whereas well-mixed populations map on to Markovian Pólya-Eggenberger urn models [62], the hierarchy of doublings in time eliminates Markovianness in the full jump-driven growth: the genetic makeup of individuals entering the population at time *t* depends on the state of the population at times of order *t*/2. Furthermore, the reinforcing influence [63] of a random draw (the size of the satellite generated by a rare long-distance jump) increases over time as *ℓ*^*d*^(*t*/2). Generalized urn-like models that incorporate these features could be useful to develop and analyze as minimal mathematical models of genetic diversity in spatially structured expanding populations. In particular, they could lead to an understanding of heterozygosity *distributions*, including limit distributions in situations where the long-time heterozygosity converges.

The coarse-grained model of blob evolution provides a route to understanding the genealogical structures left behind by jump-driven expansion, which is crucial for demographic inference. The effective population of satellites, over “generations” corresponding to doublings in time, is much simpler to describe compared to the full stochastic dynamics; analyzing the genealogical structure of this effective population in different growth regimes would be a useful first step to understanding genealogies in the full stochastic model. For instance, in the power-law growth regime (*d* < *μ* < *d* + 1), the effective population size is constant, which suggests that the genealogies of satellites are described by the Kingman coalescent under an appropriate rescaling of the time variable (essentially, log_2_(*t*) must be used in place of time for the Kingman coalescent to apply to the effective population of satellites). More generally, the temporal connection between the population at time *t* and its state at the earlier time *t*/2 is reminiscent of the mechanism underlying the seed-bank coalescent [64, 65] which might be applicable to our model in all growth regimes. Computations of genealogy statistics in the effective satellite population could be used to generate quantitative predictions for individual-based sampling statistics such as allele frequency spectra. Such predictions, which will also require evaluating the statistical relationships between samples of individuals and samples of satellites in the time-doubling hierarchy, are an exciting target for future work.

## METHODS

Simulations were implemented in the `C++` programming language. Pseudorandom numbers were generated using the Mersenne Twister engine provided in the `C++` standard library. Deme positions are quantized to an integer lattice in *d* dimensions. The simulation keeps track of all occupied demes and the allelic identity (`0` or `1`) of each deme. To avoid finite-size effects without initializing enormous arrays of mostly-empty demes, occupied demes were stored in unordered containers implemented using hash tables [66, 67] (specifically, `unordered_map` from the `C++` Standard Template Library was used). Using this approach, the effective lattice size is 2^64^ demes in 1D and 2^32^ × 2^32^ demes in 2D.

Simulations are initialized by randomly assigning allelic identities to a compact zone around the origin. In 1D, demes are filled in the range (−*N*_0_/2*, N*_0_/2], whereas in 2D all demes are filled out to a specified distance from the origin. At each simulation step, an occupied deme is chosen at random from the population as the source for a jump attempt. The jump distance, *r* is drawn at random from the probability distribution *J*(*r*) = *μr^−^*^(*μ*+1)^ (operationally, a random number *X* is drawn from the uniform real number distribution between 0 and 1, and *r* = *X^−^*^1/*μ*^ then provides a variable which follows the *J*(*r*) distribution). A random *d*-dimensional unit vector is also generated (±1 in 1D, and evenly distributed on the unit circle in 2D) and multiplied with *r*, following which each component is rounded to the nearest integer to obtain the candidate jump vector. The target deme for the jump attempt is obtained by adding the source deme position to the candidate jump vector. If the target deme is empty, it is filled with the allelic identity of the source; if it is occupied, the jump attempt is not successful.

The output of the simulations varied based on the measured quantity. For tracking the heterozygosity, it was sufficient to record the allele fractions at successive population sizes. Images of simulated populations required occupied deme positions and allelic identites to be recorded. Measurements of the MRCA positions were conducted in separate simulations in which the outbreak was begun from a single occupied deme, and a genealogical tree was maintained and updated at each successful jump. At the end of each simulation, the MRCA was recorded for pairs positioned at different center-pair distances from the origin.

Run times are determined by the number of failed attempts made as the simulation progresses towards a target population size. The run time for individual simulations ranged from a few minutes to 72h, and was significantly higher for kernels with *μ* > *d* compared to broader kernels. The upper limit was set by access to computational resources, and as a result, desired population sizes could not be achieved for the narrowest kernels. Ensemble-averaged heterozygosity measurements were obtained by averaging over 100-700 independent simulations for each set of parameters, depending on system size. MRCA distance measurements were obtained by averaging over 1000 independent simulations for *μ* = 0.6 and *μ* = 1.0, and over 400 independent simulations for *μ* = 1.4.

## Code availability

Code used to generate simulation data will be made available by the authors upon request.

## ACKNOWLEDGMENTS

Research reported in this publication was supported by a National Science Foundation Career Award (#1555330) and by a Simons Investigator award from the Simons Foundation (#327934). This research used resources of the National Energy Research Scientific Computing Center (NERSC), a U.S. Department of Energy Office of Science User Facility operated under Contract No. DE-AC02-05CH11231. This work benefited from access to the University of Oregon high performance computer, Talapas.

## Appendix A: Coarse-grained model of heterozygosity evolution

Here we develop the coarse-grained model of heterozygosity evolution based on the typical spacetime relationships between the core and satellites during jump-driven growth. The model requires prior knowledge of the function *ℓ*(*t*) specifying the growth in radius of the population core (i.e. the region surrounding the homeland which is almost fully occupied). Once *ℓ*(*t*) and a starting point *t*_0_ defining the homeland have been specified, the model allows us to generate stochastic outcomes for the evolution of neutral diversity as the population grows for times much longer than *t*_0_.

## 1. A single time-doubling “generation”

The key insight from Ref. 38 is that the typical satellite joining the core at time *t* was seeded at time of order *t*/2. Since the isolated satellite grows from a single seed through the same jump-driven process as the core, it grows up to a size *ℓ*(*t*/2), as shown schematically in main text Fig. 2**c**. The coarse-grained model is obtained by assuming that this typical behavior is followed exactly during the evolution, leading to a programmatic placement and growth of satellites around the core.

Specifically, we consider a population which has grown from a few individuals at time 0 to a known population of radius *ℓ*(*t*_0_) at time *t*_0_. Our goal is to link the neutral variation of the growing population at times much later than *t*_0_ to the state of the population at *t*_0_, which we call the initial population or homeland. We assume that alleles are uniformly distributed in the homeland; i.e. the gene pool is fully described by the fractions of various neutral alleles in the population.

In the jump-driven growth regime, the population grows through the accumulation of successive satellites, each placed at a time *t* such that it exactly touches the edge of the core population upon growing for an interval of time *t*. Given a growth function *ℓ*(*t*) for the growth of the radius of each satellite, the placement of the satellites is completely determined by *ℓ*(*t*). In 1D, this is achieved by solving for the unique satellite seeding times *τ*_*j*_ by working outwards from the core as follows:

- the first pair of satellites is seeded at the time *τ*_1_ which satisfies the equation *ℓ*(2*τ*_1_)−*ℓ*(*τ*_1_) = *ℓ*(*t*_0_), and touch the core exactly at time 2*τ*_1_ (*τ*_1_ satisfying the relation must be found numerically, and has a value *τ*_1_ ≳ *t*_0_/2);
- the following pairs are seeded at *τ*_*j*_ such that *ℓ*(2*τ*_*j*_) − *ℓ*(*τ*_*j*_) = *ℓ*(2*τ*_*j*−1_)+*ℓ*(*τ*_*j*−1_), lined up to touch the previously seeded satellites exactly at time 2*τ*_*j*_.

This protocol can be generalized to 2D, where isotropic expansion of the core requires the addition of roughly *πℓ*(2*τ*_*j*_)/*ℓ*(*τ*_*j*_) identically-sized satellites to encircle the population (rather than a pair of satellites of size *ℓ*(*τ*_*j*_), one on either side, for the 1D population). In both cases, the time-doubling hierarchy of key jumps dictates that the homeland is able to seed satellites at successive times *τ*_*j*_, found numerically through the above protocol, until the value of *τ*_*j*_ crosses *t*_0_. The furthest satellites seeded in this manner therefore take root at times very close to *t*_0_, and at distances of order *ℓ*(2*t*_0_) from the core. They grow to reach a size close to *ℓ*(*t*_0_), before they run into their adjacent satellites. The expansion of the population from radius *ℓ*(*t*_0_) to *ℓ*(2*t*_0_) through the accumulation of satellites is considered the first “time-doubling generation” in the model. The number of satellites generated through the above protocol is called *g*_1_ for the first doubling (and *g*_*m*_ for successive doublings indexed by *m*). Main text Fig. 5 shows examples of satellites generated from the algorithm over successive time-doublings.

Although the placement and size of satellites is completely deterministic in the coarse-grained model, stochasticity is maintained in the random choice of allelic identity of each satellite from the core population. First we consider the simplest case of two alleles `A` and `B`, with initial fractions *p*_0_ and 1 − *p*_0_ in the homeland respectively. The allelic identities of satellites are then chosen as random draws with these probabilities. As a result, the final fraction *p*_1_ of allele `A` in the time-doubled population is a random variable. From the examples in main text Fig. 5, it is apparent that the allele fractions of the population after the time-doubling depend on the sizes of the added satellites. The population of the *i*th satellite os *ω*_*d*_*ℓ*(*t*_1*,i*_), where *ω*_1_ = 2, *ω*_2_ = *π*; *i* ∈ {*1*, …, *g*_1_} indexes the satellites; *t*_1,*i*_ is the growth time of satellite *i* in the 1st doubling; and *i* = 0 refers to the homeland (so *t*_1,0_ ≡ *t*_0_). To find the expectation value of *p*_1_ across independent stochastic doublings, we note that the probability of a random draw from the population being allele `A` is the weighted probability of a draw from the homeland, or from each satellite, being allele `A`. For the biallelic homeland, this probability is *p*_0_; for each monoallelic satellite, it equals the probability of the satellite having identity `A`, which is also *p*_0_. As a result,

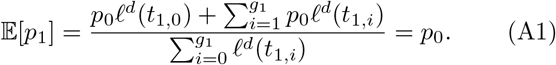

Therefore, the allele fraction is conserved *on average* during the time-doubling generation. Note that this result also holds for a population of *M* distinct neutral alleles in the gene pool: for each allele, the population can still be divided into a fraction *p*_0_ of that allele and 1 − *p*_0_ for all other alleles, and the expectation value of each individual allele fraction is preserved by the random draws.

Whereas the averages of the allelic fractions do not change, we expect their variances to evolve with time-doubling generations. A single measure related to the variances of allele frequencies is the heterozygosity, defined as the probability that a random sample of two individuals from the population is biallelic. For a population of *M* distinct neutral alleles with fractions *p*_0,1_, *p*_0,2_, …, *p*_0*,M*_, the heterozygosity is

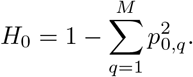

We now aim to relate the expected final heterozygosity *H*_1_ of the population after the first time doubling, in terms of *H*_0_. We can break down the probability of a sampled pair in the time-doubled population being biallelic in terms of the origin of the two samples:

- 0 if both samples were drawn from the same new satellite;
- *H*_0_ if both samples were drawn from the homeland;
- *H*_0_ if one sample was from a homeland and one from a new satellite;
- *H*_0_ if both samples were drawn from two different new satellites.

(The last two are true because the distribution of types of the new satellites is exactly the same as the distribution of individuals in the homeland.) The probability of choosing from the *i*th satellite twice in two independent draws is 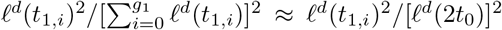, where we use the fact that the doubling protocol places the outermost satellites at a distance close to *ℓ*(2*t*_0_) from the origin, so the total population after the doubling is 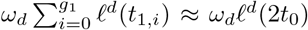. Therefore the expected heterozygosity after the doubling is

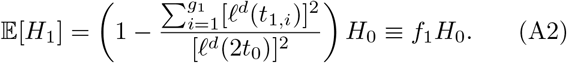

We note that the above result is independent of the number of distinct alleles, as it is expressed purely in terms of the heterozygosity. Therefore, while our simulation study is restricted to biallelic simulations, our results for the evolution of neutral heterozygosity based on the coarsegrained model apply more generally.

## 2. Successive time-doublings

The above protocol generates a population of radius ≈ *ℓ*(2*t*_0_) from the gene pool at time *t*_0_. To continue the time-evolution, the collection of deterministically-placed satellites with stochastic allelic identities is treated as a new *homogeneous* homeland with allele fractions *p*_1,1_, *p*_1,2_, …, *p*_1*,M*_, and size *ℓ*(2*t*_0_). This homogenized population is used to generate a set of *g*_2_ satellites, taking the population size to roughly *ℓ*(4*t*_0_), which comprises the next generation. The homogenization is repeated for subsequent time-doubling generations to evaluate the long-time behaviour of the range expansion. In general, the *m*th doubling expands the population from radius *ℓ*(2^*m−*1^*t*_0_) to *ℓ*(2^*m*^*t*_0_), through *i* ∈ {1, 2, …, *g*_*m*_ new satellites with radii *ℓ*^*d*^(*t*_*m,i*_) determined by the protocol described above. The source size of the *m*th doubling is *ℓ*(*t*_*m,*0_) = *ℓ*(2^*m−*1^*t*_0_).

Generalizing Eq. (A1) over successive time doublings (indexed by *m*) for any given allele (indexed by *q*), the expectation value 𝔼[*p*_*m,q*_] simply equals the previous value *p*_*m*−*1,q*_. Therefore the sequence of random variables *p*_*m,q*_ for successive values of *m* comprises a martingale. Since allele fractions are bounded to take on values between 0 and 1 inclusive, *p*_*m,q*_ has a random limit by the martingale convergence property. This convergence is independently achieved for each allele; i.e. over infinitely many generations, each allele fraction approaches a random limit, with the constraint that the allele fractions sum to one.

By using the random limit property, we can heuristically anticipate the outcomes for heterozygosity at long times. Focusing on the fraction of a particular allele in the population, the existence of a limit states that at upon carrying out the time-doublings infinitely many times, fluctuations in the allele fraction due to random sampling have died out completely. This can happen in two ways. One possibility is that the number of satellites generated has become infinitely large, which implies that *g*_*m*_ must grow between successive doublings. In this case, the limiting fraction could take on values between 0 and 1, allowing for heterozygosity to be preserved (although it is not guaranteed to be preserved: lim_*m*→∞_ g_*m*_ = ∞ is a necessary, but not sufficient, condition for the heterozygosity to converge to a non-zero value). Alternatively, if the number of satellites generated does not grow but rather stays finite, fluctuations in the allele fraction can only be avoided if the allele fraction approaches 0 or 1; i.e. heterozygosity must be completely lost.

To quantitatively analyze these trends, we compute the recursion for the expected heterozygosity over *n* time-doublings,

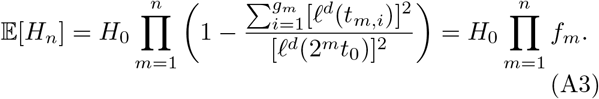

Note that Eq. (A3) contains only the radii *ℓ*^*d*^(*t*_*m,i*_) and thus depends on the deterministic part of the update protocol. By following the protocol described above to generate the precise satellite sizes, the values *f*_*m*_ can be numerically generated for different growth rules *ℓ*(*t*) and homeland sizes *t*_0_.

## 3. Approximating the heterozygosity factors

Since the convergence of the product in Eq. (A3) depends on the behavior of the heterozygosity factors *f*_*m*_ with increasing *m*, we now analyze how *f*_*m*_ depends on *m* for the various asymptotic growth forms summarized in Table 1.

At the *m*th doubling, satellites are generated to fill up a region of space extending out to *ℓ*(2^*m*^*t*_0_), and the largest satellite generated has radius approximately *ℓ*(2^*m−*1^*t*_0_). We expect these two length scales to dictate the behavior of the sum 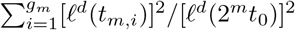 in *f_m_*, up to geometric factors. A crude estimate is obtained by isolating the effect of the growth in the number of satellites as the doublings proceed, and ignoring variations in the satellite size. Assuming that each satellite in the *m*th doubling has radius equal to the average radius 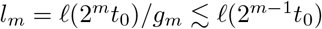, we estimate the heterozy-gosity factors as

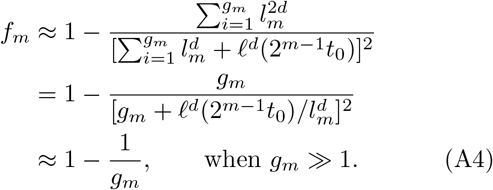

Geometrically, the number of satellites scales with the ratio [*ℓ*(2^*m*^*t*_0_)/*ℓ*(2^*m−*1^*t*_0_)]^*d*^, (strictly, a lower bound to *g_m_*). By absorbing corrections to this form into an overall constant, we hypothesize the following approximation in terms of the growth function *ℓ*(*t*):

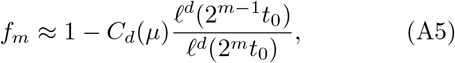

where the constant *C_d_*(*μ*) captures geometric effects stemming from the variation in satellite sizes, which are ignored in Eq. (A4).

We test our assumptions by comparing the approximations, Eq. (A4) and Eq. (A5), with results of the satellite placement protocol described above from which the satellite counts *g*_*m*_ and heterozygosity factors *f*_*m*_ are computed numerically. Results are shown in Fig. A1 In all regimes of the asymptotic growth rules *ℓ*(*t*), we find that Eq. (A5) successfully captures the results of the numerically-evaluated heterozygosity evolution for a range of homeland sizes, with constants *C*_*d*_(*μ*) of order one. Interestingly, the second approximation, Eq. (A5), performs remarkably well in capturing trends in heterozygosity factors, even when Eq. (A4) does not perform as well in reproducing the number of satellites generated upon doublings. In essence, Eq. (A5) makes two quantitative errors in estimating the sum 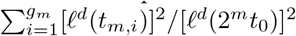: it replaces the number of terms in the sum with [*ℓ*(2^*m*^*t*_0_)/*ℓ*(2^*m−*1^*t*_0_)]^*d*^ which is a lower bound to the true number *g*_*m*_ and hence underestimates it; it also replaces the sizes of satellites in the numerator with *ℓ*^*d*^(2^*m−*1^*t*_0_), which provides an upper bound to the size of the new satellites and hence overestimates it. These opposing contributions appear to perform better than Eq. (A4), which has the correct number of satellites but ignores variations in the size of satellites.

**FIG. A1.**
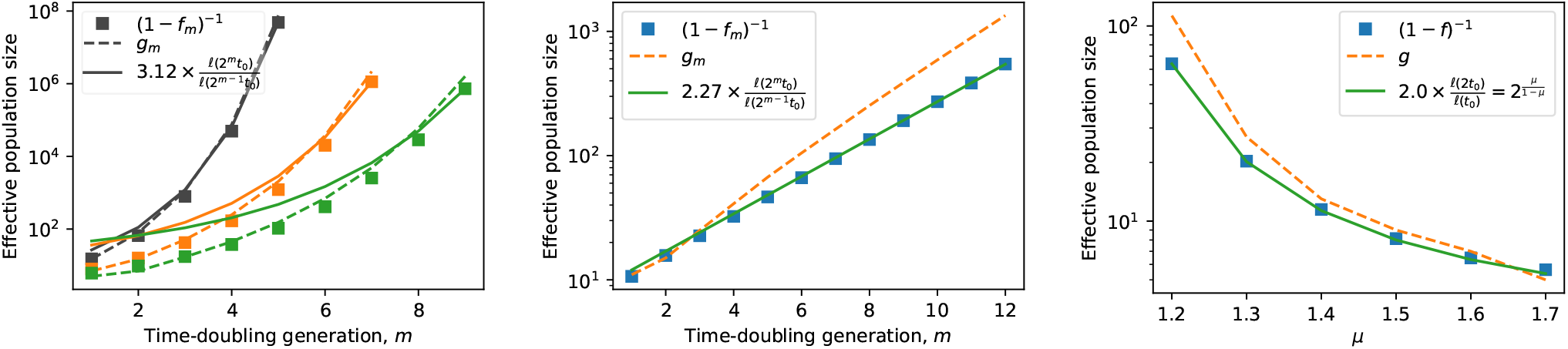
Test of approximate form for heterozygosity factors *f*_*m*_. Graphs show the effective population size of satellites 1/(1 − *f*_*m*_) implied by the growth factors *f*_*m*_ defined in Eq. (A3), with satellite growth times computed from the algorithm specified in Appendix A for a 1D expansion (symbols). The algorithm is begun with a homeland size of *N*_0_ = 100. The *ℓ*(*t*) functions used are the asymptotic forms from Table I (main text). Also shown are two estimates of the effective population size: the number of satellites *g*_*m*_ generated by the algorithm in the *m*th doubling (dashed line), and the ratio *ℓ*(2^*m*^*t*_0_)/*ℓ*(2^*m*−1^*t*_0_) multiplied by a constant obtained from a least-squares fit (solid line). **a**, Kernels with (from left to right) *μ* = 0.2, 0.4, 0.5 respectively. The effective population size grows faster than exponentially with the number of time-doubling generations. **b**, *μ* = 1. The effective population size grows exponentially with time-doublings; the growth in number of satellites generated overestimates the growth in the effective population size computed from *f_m_* whereas Eq. (A5) is more accurate. **c**, Kernels with 1 < *μ* < 2. The scale-free nature of the power-law growth form in this regime ensures that *f* depends only on *μ* and does not change with successive time-doublings. The effective population size as a function of *μ* is well-reproduced by both approximations, with Eq. (A5) proving more accurate.

## 4. Convergence properties

We now justify the use of our approximate forms above, and in particular the behavior of the size ratios *ℓ*(2^*m−*1^*t*_0_), in drawing conclusions about the convergence properties of the heterozygosity over many time-doubling generations. Convergence of the product in Eq. (A3) as *n* → ∞ can be recast as a convergence of the sum of logarithms of the terms, which in turn requires convergence of the sum

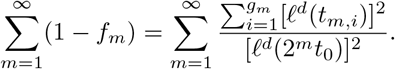

Qualitative aspects of the convergence of this sum are determined by the rate of change (or absence thereof) of successive terms, and are unaffected by overall constant factors. The approximations we have introduced, Eq. (A4) and Eq. (A5), correctly capture the scaling of successive terms in the sum with the time-doubling generation *m*, and therefore provide the correct qualitative behavior of the heterozygosity in the coarse-grained model in different growth regimes. In the following section, we show that the approximations can also reproduce many features of the heterozygosity evolution in the lattice-based simulations.

## Appendix B: Quantitative predictions for simulation results

In the previous section, we developed approximate expressions for the evolution of heterozygosity over successive “time-doubling generations” in a coarse-grained model for jump-driven growth. Analysis of the coarsegrained model showed qualitative differences for the fate of heterozygosity in the different growth. In the main text, we showed that these qualitative differences are also present in the fully stochastic jump-driven range expansions, which were simulated using the lattice-based model. Here, we develop additional predictions for the dependence of the heterozygosity on homeland size and real time using the coarse-grained model, and compare these predictions to ensemble-averaged quantities measured from the stochastic simulations. We find that Eq. (2), combined with the geometrically-motivated scaling of *f_m_*, can capture the overall scaling of the final heterozygosity with homeland size and kernel exponent in simulations. In the marginal and stretched-exponential growth regimes (*μ* ≤ *d*), which show the strongest separation of scales between time-doublings underlying the model, near-quantitative agreement is achieved through the introduction of a minimal number of fit parameters.

## 1. Stretched-exponential growth (0 < *μ* < *d*)

Following the approximations detailed above, we have 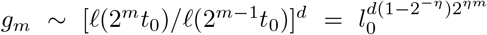. For this functional form, no closed form exists for the *n* → ∞ limit of the product in Eq. (2). However, the first several terms dominate due to the extremely fast increase in *g*_*m*_ with *m*, and a rough approximation to the deviation of the final heterozygosity from *H*_0_ is obtained by keeping only the most significant term:

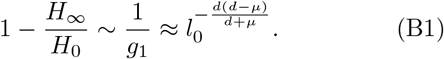

**FIG. A2.**
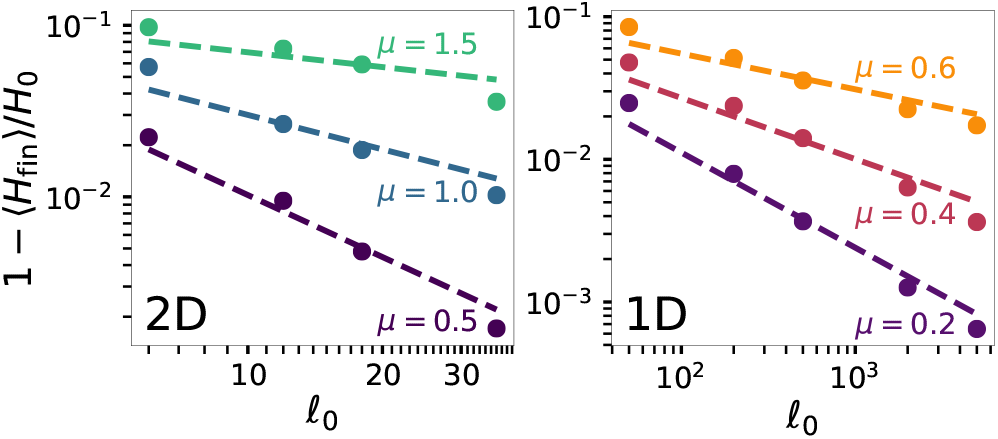
Homeland-size dependence of average final heterozygosity for *μ* < *d*. Symbols show the fractional deviation of the final heterozygosity from its initial value for simulations of both planar (left) and linear (right) habitats. Data are plotted against the homeland radius 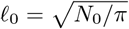 in 2D and *ℓ*_0_ = *N*_0_/2 in 1D. The final population size is *N* = 10^7^, large enough for the heterozygosity to have converged to its constant value in all cases. Dashed lines show the relation 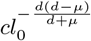 from Eq. (B1), with the magnitude parameter *c* determined by a fit to the data points for each kernel (fit value varies between 0.13 and 0.16 for 2D curves, and between 0.17 and 0.24 for 1D).

Up to an overall multiplicative factor for each kernel, Eq. (B1) captures much of the variation in average heterozygosity values as a function of homeland size in simulations despite the approximations involved, see Fig. A2. In particular, the approximation correctly reproduces the reduction in sensitivity of the final heterozygosity to homeland size as *μ* is increased.

## 2. Marginal growth (*μ* = *d*)

The scaling of the number of satellites at doubling *m* suggests the following crude approximation to the heterozygosity reduction factors determined via geometry in Eq. (2):

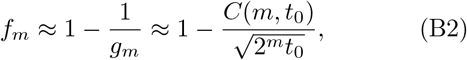

where *C*(*m*, *t*_0_) captures the effect of geometrically-determined size variations in satellites at the *m*th doubling. The residual variation in *f*_*m*_ due to *C*(*m*, *t*_0_) is small relative to the exponential dependence, so we may ignore this variation and treat *C*(*m*, *t*_0_) = *C*_*d*_(*t*_0_) as a constant during growth from the homeland size *t*_0_. In this case, *f*_*m*_ approaches one fast enough for the product in Eq. (2) to converge to a finite value as *n* → ∞. Therefore, the deterministic model predicts that in the marginal case, neutral heterozygosity is partially preserved, converging to a finite value 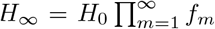 which lies between *H*_0_ and zero. In contrast to the power-law growth regime, genetic drift is indeed mitigated by growth when *μ* = *d*: the growth in the *effective* population size is fast enough to overcome drift.

**FIG. A3.**
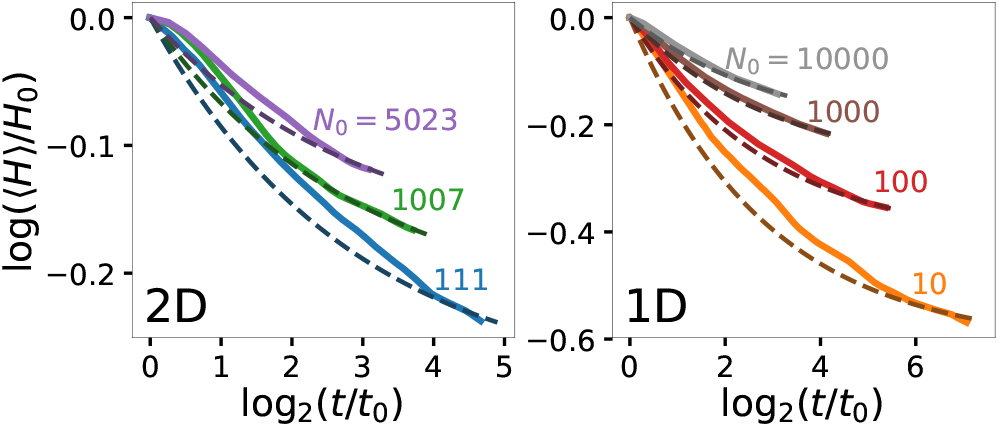
Decay of heterozygosity during marginal growth, *μ* = *d*. Solid lines show the decay of average heterozygosity for different homeland sizes (labels) with *μ* = 2 in 2D (left) and *μ* = 1 in 1D (right). Data are shown as a function of doublings in time, where *t*_0_ = *ℓ*^−1^(*ℓ*_0_) is computed from the initial homeland radius *ℓ*_0_. Dashed lines show fits to the approximate expression from the coarse-grained model, Eq. (B3), with *C*_*d*_ as a fit parameter. Fit values for individual *N*_0_ curves are (*N*_0_ = 111: *C*_*d*_ = 1.015); (1007: 1.109); (5023: 1.082); (10: 0.730); (100: 0.901); (1000: 0.933); (10000: 1.008).

At first glance, the simulated evolution of average heterozygosity for *μ* = *d* does not show convergence up to system sizes of *N* = 10^8^ (Fig. 4). However, this observation is not inconsistent with the conclusions from the coarse-grained model: the convergence of the product in Eq. (2) occurs over many “generations” constituting doublings in time *t*, which correspond to population growth ~ *ℓ*^*d*^(*t*) spanning many orders of magnitude. Since *g*_*m*_ ≫ 1, log *f*_*m*_ can be approximated as 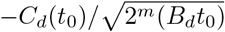, and the product in Eq. (2) is exactly evaluated to give

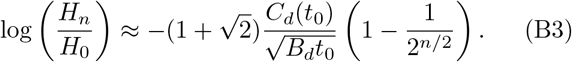

We find that Eq. (B3) reproduces the heterozygosity evolution measured in simulations, with fit values *C*_*d*_(*t*_0_) of order one (Fig. A3). In all cases, the populations are several doublings away from converging to the predicted final value, which would only be observable in populations that are larger by many more orders of magnitude.

## 3. Power-law growth (*d* < *μ* < *d* + 1)

In the main text, we argued that the scale-free nature of power-law growth functions leads to heterozygosity factors that remain constant over successive doublings in the deterministic model. Constant heterozygosity factors in Eq. (A3) imply an exponential decay of heterozygosity with the “time-doubling generations” *n* ≈ log_2_(*t/t*_0_). However, in real time, the heterozygosity experiences a much gentler power-law falloff: 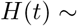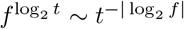. This form can be recast as a function of population size using *N ~ t^d/^*^(*μ*−*d*)^, to obtain

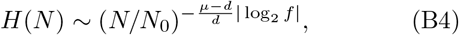

with a rough estimate for *f* provided by the expected number of satellites as log_2_ *f* ≈ log_2_(1 − 1*/N*_e_) ≈ −2^*−d*/(*μ*−*d*)^/ log 2. This estimate is compared to the results of simulations in Fig. A4. The simulations are consistent with a power-law decay of average heterozygosity with population size, although computational limitations restrict the observable heterozygosity range. The rough estimate of the decay exponent qualitatively reflects the variation in the decay with kernel exponent, but it is quantitatively inaccurate, especially in *d* = 2. This discrepancy likely reflects the inadequacy of the asymptotic growth rules to describe the intermediate growth periods relevant to simulations [38], and the fact that the separation between populations at times *t* and 2*t* is much weaker for power-law growth compared to the other regimes of jump-driven growth.

**FIG. A4.**
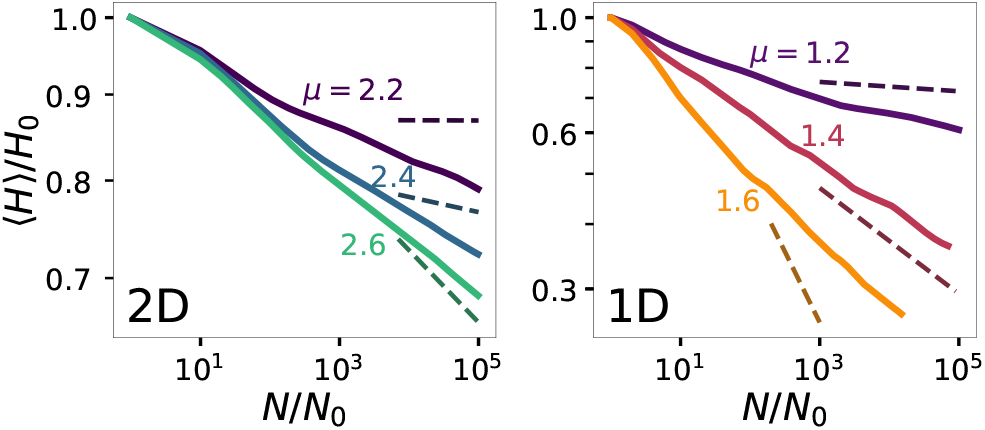
Decay of average heterozygosity in the power-law growth regime, *d* < *μ* < *d* + 1. Solid lines show the decay of average heterozygosity for different kernels (labeled) in the regime where the asymptotic growth rule is power-law with time. Homeland size is *N*_0_ = 1007 for all kernels in 2D (left), and *N*_0_ = 1000 for all kernels in 1D (right). Dashed lines indicate power-law decay with exponent −2^−*d*/(*μ*−*d*)^(*μ − d*)/(*d*log 2) from the rough estimate motivated by the deterministic model.

## Appendix C: Robustness of observed patterns to model variations

Although simulation results in the main text were reported for well-mixed homelands with two alleles in equal proportion, we show here that our conclusions regarding the breakup of sectors and the distinct regimes of heterozygosity decay are robust to variations in the model.

## 1. Variations in homeland structure

Homelands with a patchy distribution of neutral alleles (i.e. individuals are geographically segregated by allelic identity in the initial population) might be expected to favor sectors during the range expansion. To test the effect of such structured homelands on subsequent patterns, we performed range expansions simulations with *q* = 4 distinct alleles in a large sectored homeland with *N*_0_ = 6074, as shown in Fig. A5. Although independent trials were initialized with the same homeland, stochastic duplication and migration events led to variations in the final outcome. We found that the sectoring of the homeland persisted (up to early stochastic fluctuations) for *μ* = 4, but was erased for jump-driven growth in both the power-law (*μ* = 2.6) and stretched-exponential (*μ* = 1.5) regimes (Fig. A5. This observation is consistent with the expansion mechanism. When *μ* < *d* + 1, satellite clusters are seeded by jumps whose length is many times the size of the core population. As a result, satellites do not have a strong preference for landing in the angular sector that extends radially outward from any one patch, and sectors are broken up into blobs and speckles within a few time-doubling generations. Consequently, the specific structure of the homeland does not leave a significant impact on the asymptotic heterozygosity evolution at long times. This feature is built into the coarse-grained model, which treats the population between successive time-doublings as well-mixed without regard to the spatial distribution of alleles.

Since the qualitative features of the patterns persist for sectored homelands, the trends in heterozygosity evolution are reproduced as well: heterozygosity is preserved for *μ* = 4 and *μ* = 1.5, but decays inexorably as the expansion progresses when *μ* = 2.6. Compared to a well-mixed homeland, the sectored homeland favors the preservation of heterozygosity in the constant-speed expansion (*μ* = 4) by seeding sectors with a similar pattern to the homeland. By contrast, the heterozygosity trend is largely unaffected by the homeland structure in the jump-driven growth regime. Simulations over a wider range of kernels for a smaller homeland (*N*_0_ = 111) with the same angular distribution of alleles confirmed that the drop in heterozygosity for *d* < *μ* < *d* + 1 persists for sectored homelands (Fig. A6(a)).

We also measured heterozygosity evolution in expansions from homelands in which each lattice site was assigned a distinct allelic identity, corresponding to an initial heterozygosity of one. The resulting heterozygosity trend (Fig. A6(b)) is similar to that reported for the two-allele simulations in the main text, confirming that the results are independent of the precise allelic composition of the homeland.

**FIG. A5.**
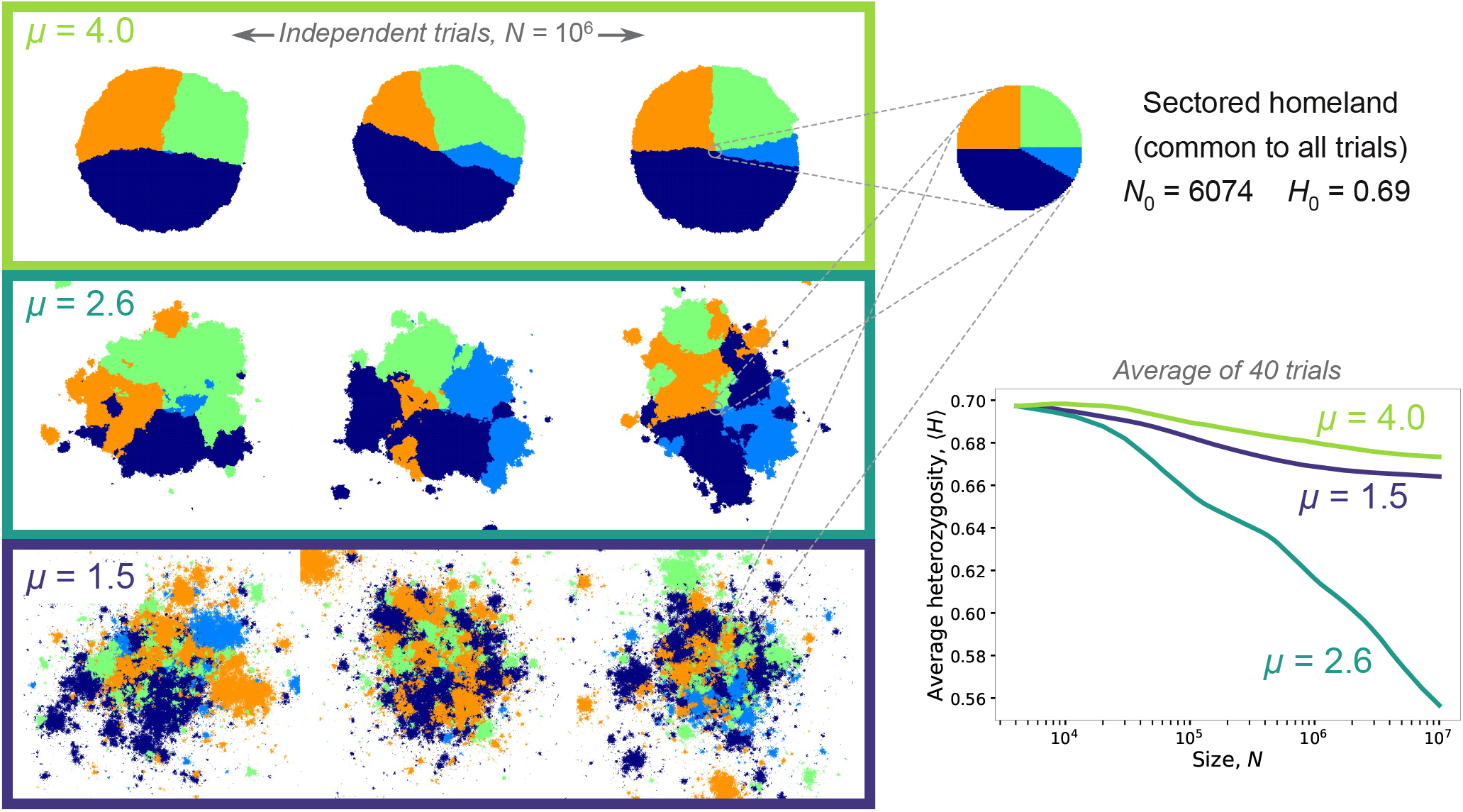
Blobs and speckles persist for structured homelands. Left, a sampling of snapshots at *N* = 10^6^ of independent range expansions from a homeland made up of four monoallelic sectors of fractions 1/4, 1/4, 5/12, and 1/12. The homeland (shown at top right) is shared among all trials and kernels. The choice of homeland encourages the formation of sectors with a similar pattern for *μ* = 4, although the angular distribution of alleles is not perfectly maintained due to early stochastic fluctuations. However, blobs and speckles are still generated in the jump-driven regime, *μ* = 2.6 and *μ* = 1.5. The average heterozygosity measured from 40 independent trials (bottom right) follows the trends expected from the coarse-grained model: as the expansion progresses, diversity is lost for *μ* = 2.6 which is in the power-law growth regime, whereas diversity persists in the other growth regimes.

**FIG. A6.**
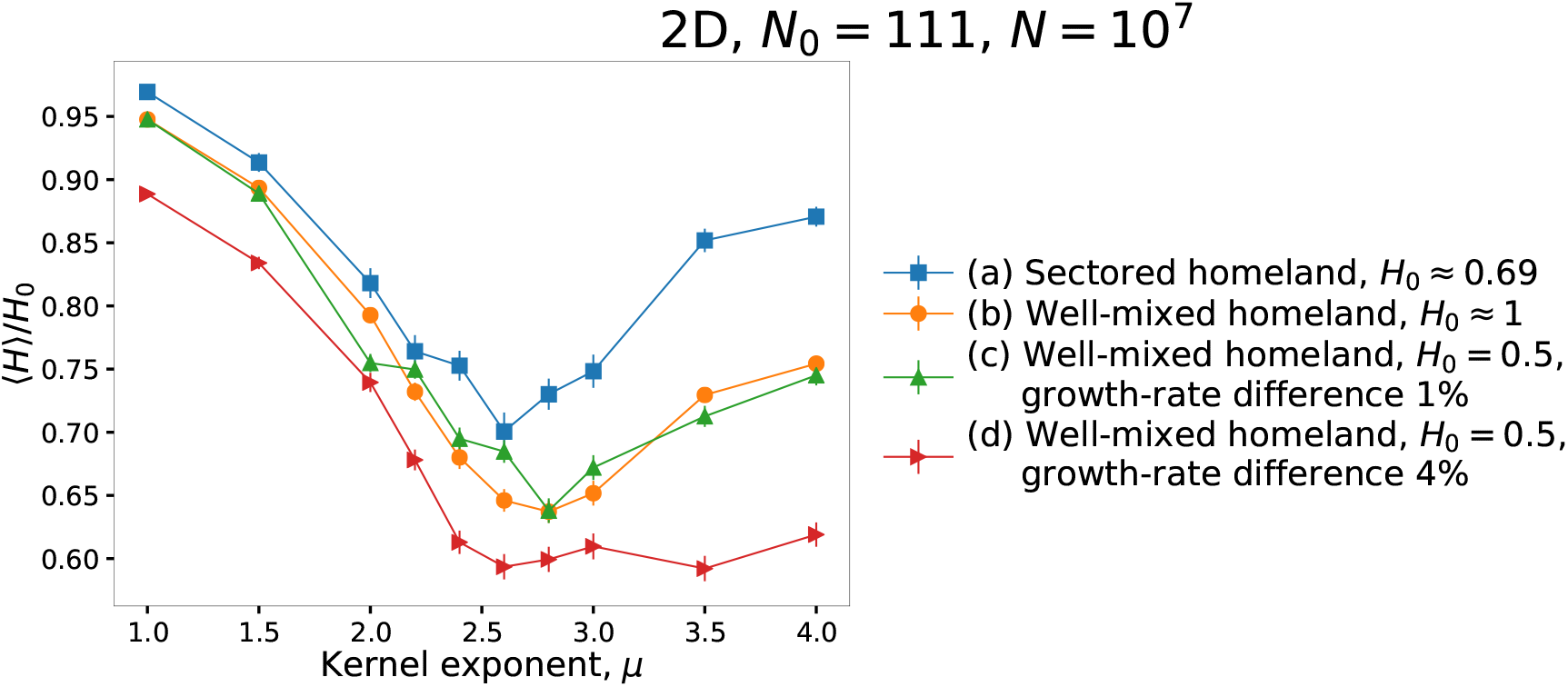
Heterozygosity decay is robust against model variation. Average final heterozygosity at *N* = 10^7^ for expansions from homelands with *N*_0_ = 111, with various modifications of the initial or growth conditions. (a) Sectored homeland with four distinct alleles, patterned as shown in Fig. A5 but with a smaller radius. (b) Well-mixed homeland with *N*_0_ distinct alleles, i.e. each individual in the homeland has a unique allelic identity. (c) Well-mixed homeland with a mix of two alleles in equal proportion, where the duplication rate of one allele is 1% higher than the other. (d) As in (c), with 4% higher duplication rate for one allele.

## 2. Fitness variations

Real populations are likely to harbor slight fitness variations among different alleles. Over long times, fitness variations lead to a decay of heterozygosity during the expansion even for well-mixed populations, as the fittest alleles increase their representation in the population. Similarly, for populations expanding through the advance of a constant-speed front, sectors of fitter alleles grow faster than less-fit alleles and eventually dominate the range expansion. Therefore, in the presence of fitness differences, we expect population-level heterozygosity to decay to zero at long times for all growth regimes. However, when variations in fitness are small, spatial effects would still be apparent at intermediate times. A thorough analysis of the effect of fitness variations on the dispersal-driven range expansion model would be interesting to pursue, but is beyond the scope of the present work.

Here, we show that moderate fitness variations do not immediately erase the heterozygosity decay expected in the power-law growth regime for the population sizes simulated in the main text. We ran simulations starting from well-mixed homelands with an equal mix of two alleles as in the main text, but set the duplication rate of demes belonging to one allele to be higher by a small fraction. When the growth rate difference between the alleles was set to 1%, the average heterozygosity at a population size of 10^7^ remained markedly lower for the power-law growth regime *d* < *μ* < *d* + 1 compared to the other two regimes, see Fig. A6(c). To bring down the average heterozygosity in the constant-speed expansion regime *μ* > *d* + 1 to similarly low values at *N* = 10^7^, the fitness variation needs to be increased to 4% (Fig. A6(d)). These results show that the competition between coarsening and diversification during jump-driven range expansions persists for moderate fitness variations among alleles.

**FIG. A7.**
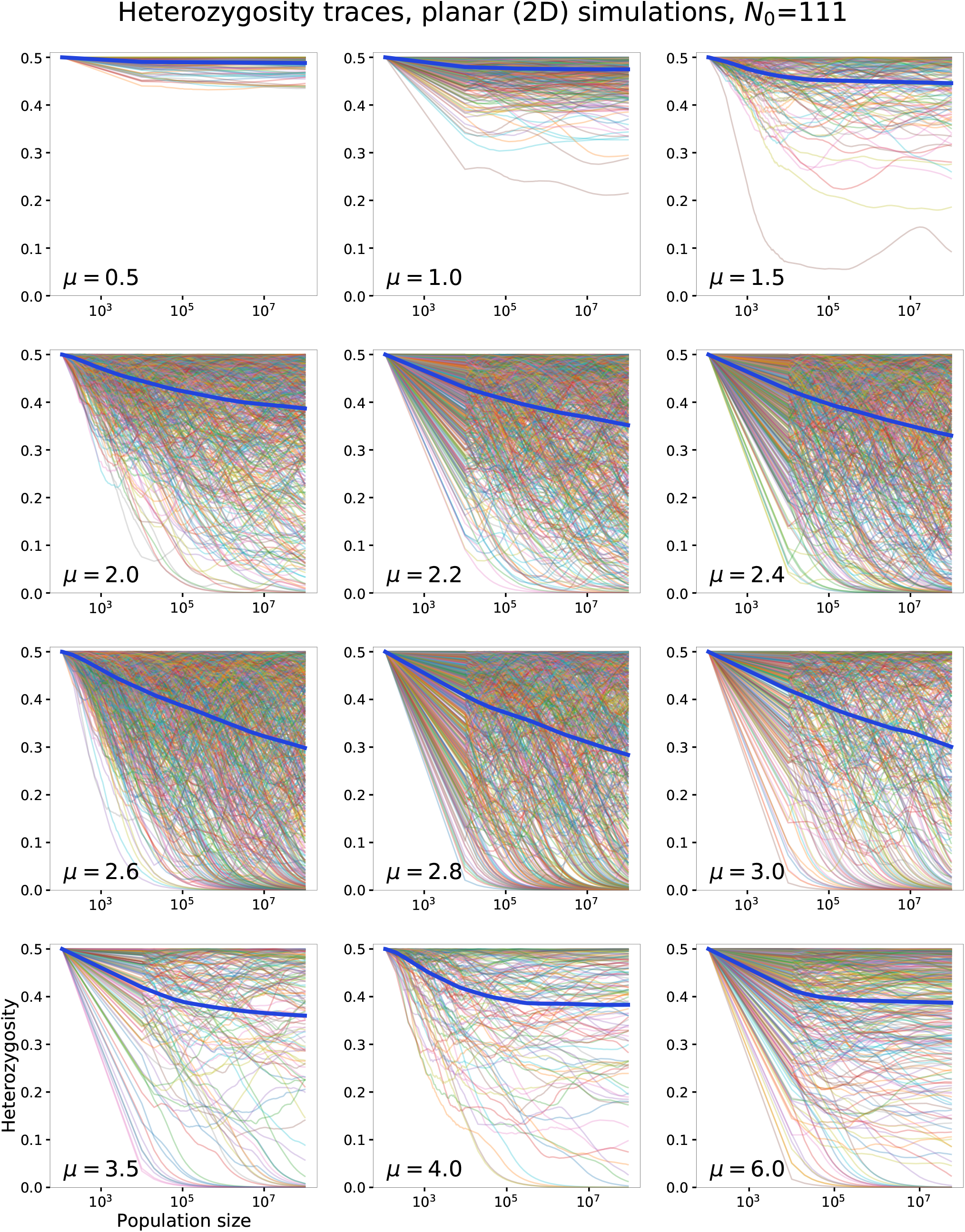
Heterozygosity evolution for 2D simulations. Panels show the heterozygosity from individual simulations (thin lines) and the ensemble-averaged heterozygosity (thick line) for all kernels reported in Fig.4a of the main text.

**FIG. A8.**
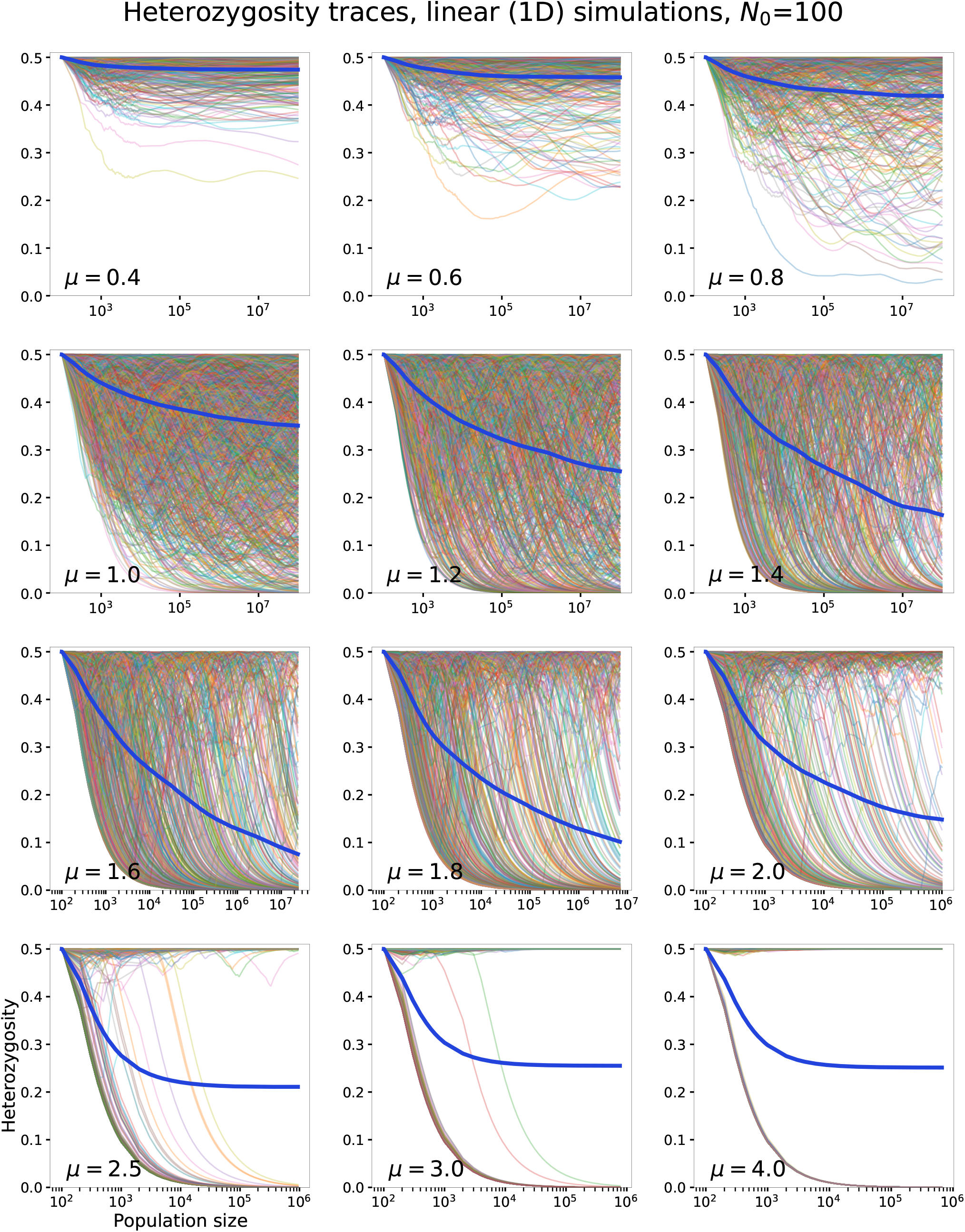
Heterozygosity evolution for 1D simulations. Panels show the heterozygosity from individual simulations (thin lines) and the ensemble-averaged heterozygosity (thick line) for all kernels reported in Fig.4b of the main text.

